# A human stomach cell type transcriptome atlas

**DOI:** 10.1101/2023.01.10.520700

**Authors:** S Öling, E Struck, MN Thorsen, M Zwahlen, K von Feilitzen, J Odeberg, F Pontén, C Lindskog, M Uhlén, P Dusart, LM Butler

## Abstract

The identification of cell type-specific genes and their modification under different conditions is central to our understanding of human health and disease. The stomach, a hollow organ in the upper gastrointestinal tract, provides an acidic environment that contributes to microbial defence and facilitates the activity of secreted digestive enzymes to process food and nutrients into chyme. In contrast to other sections of the gastrointestinal tract, detailed descriptions of cell type gene enrichment profiles in the stomach are absent from the major single cell sequencing-based atlases. Here, we use an integrative correlation analysis method to predict human stomach cell type transcriptome signatures using unfractionated stomach RNAseq data from 359 individuals. We profile parietal, chief, gastric mucous, gastric enteroendocrine, mitotic, endothelial, fibroblast, macrophage, neutrophil, T-cell and plasma cells, identifying over 1600 cell type-enriched genes. We uncover the cell type expression profile of several non-coding genes strongly associated with the progression of gastric cancer and, using a sex-based subset analysis, uncover a panel of male-only chief cell-enriched genes. This study provides a roadmap to further understand human stomach biology.

## INTRODUCTION

The gastrointestinal (GI) tract is a multiple organ system which can be divided into upper and lower parts, the physical properties and cellular characteristics of which reflect their different roles in digestion, absorption of nutrients, and excretion of waste products (de Santa Barbara, van den Brink and Roberts, 2003; Choi *et al*., 2014; Thompson, DeLaForest and Battle, 2018). The stomach, a hollow muscular organ in the upper GI tract, produces an array of acids and gastric enzymes, acting as a reservoir for the mechanical and chemical digestion of ingested food (Kim and Shivdasani, 2016). The constituent cell types of the stomach include parietal cells, chief cells, gastric mucous cells, gastric enteroendocrine cells, mitotic cells, endothelial cells, fibroblasts, and various immune cells (Gremel *et al*., 2015; Uhlen *et al*., 2015). In contrast to lower sections of the GI tract, descriptions of the cellular transcriptional landscape in the stomach are lacking, with this organ absent from large scale single cell sequencing (scRNAseq) initiatives, such as Tabula Sapiens (Tabula Sapiens *et al*., 2022) and the Human Cell Atlas (Regev *et al*., 2017). Where scRNAseq has been used to profile gene expression in the adult stomach, studies have typically focused on specific cell types, such as the epithelia (Busslinger *et al*., 2021; Tsubosaka *et al*., 2022), or in pathological states such as gastric cancer (P. Zhang *et al*., 2019; Sathe *et al*., 2020; Wang *et al*., 2021; Kim *et al*., 2022). Whilst scRNAseq studies provide high resolution of individual cell (sub)type gene expression profiles, challenges remain, including artefactual modification of gene expression due to cell removal and processing (O’Flanagan *et al*., 2019; Denisenko *et al*., 2020; Massoni-Badosa *et al*., 2020), compromised read depth, and difficulties with data interpretation (Gawad, Koh and Quake, 2016; Jiang *et al*., 2022). As a limited number of biological replicates are typically analysed, underestimation of biological variance can increase the likelihood of potential false discoveries (Squair *et al*., 2021; Denninger *et al*., 2022).

Non-coding RNA is emerging as a novel, important class of molecules, involved in the maintenance of healthy stomach tissue, and the development and progression of gastric cancer (Gao *et al*., 2020; Razavi and Katanforosh, 2022), but to date there is no overall description of stomach cell type enriched non-coding RNAs.

Here, we analysed 359 bulk RNAseq human stomach samples to identify over 1600 genes with cell type-enriched expression, using our previously developed integrative correlation analysis (Butler *et al*., 2016; Dusart *et al*., 2019; Norreen-Thorsen *et al*., 2022). Gastric mucous cells had the highest number of predicted protein-coding and non-coding enriched transcripts and represented the primary site of expression of genes that were tissue enriched in stomach over other tissue types. Gastric enteroendocrine cells expressed a panel of non-coding transcripts that are also selectively expressed in pancreatic and intestinal endocrine cells, indicting a common function in these cell types. Several of the identified cell type enriched non-coding genes have previously been associated with the progression of gastric cancer, but until now the cell type site of expression had not been described. Sex subset analysis revealed a high global similarity in cell type transcriptomes between males and females, but a panel of chief cell enriched Y-linked genes were identified. Data is available through the Human Protein Atlas (HPA) portal (www.proteinatlas.org/humanproteome/tissue+cell+type/stomach).

## RESULTS

### Identification of cell type transcriptome profiles in stomach

#### Cell type reference transcripts correlate across unfractionated RNAseq data

To identify stomach cell type-enriched transcriptome profiles, we conducted an analysis based on our previously developed method (Butler *et al*., 2016; Dusart *et al*., 2019; Norreen-Thorsen *et al*., 2022), using human stomach bulk RNAseq data (N=359) from Genotype-Tissue Expression (GTEx) portal V8 (https://gtexportal.org) (Consortium, 2015). Each sample was unfractionated and thus contained a mix of cell types (Figure 1 A.i), which contribute differing proportions of transcripts that are measured by RNAseq (Figure 1 A.ii). For each major constituent stomach cell type, candidate ’virtual markers’ (’reference transcripts’ [*Ref*.*T*.]) were selected based on: (**i**) older ’none-omics’ studies (Hassan, Toor and Ahmad, 2010) (**ii**) in-house proteomic profiling of stomach tissue (Gremel *et al*., 2015; Uhlen *et al*., 2015), (**iii**) scRNAseq data (Busslinger *et al*., 2021; Karlsson *et al*., 2021) and (**iv**) databases collated from multiple sources, e.g. <u>Cell Marker</u> (X. Zhang *et al*., 2019) and <u>PanglaoDB</u>(Franzen, Gan and Bjorkegren, 2019) (Figure 1B). Three markers were slected for each cell type, based on the following criteria: (**i**) A high corr. (>0.85) between *Ref*.*T*. <u>within</u> each cell type panel (Figure 1 C and Table S1, Tab 1), indicating *cell type co-expression*: parietal cells (PAC) [*ATP4B, MFSD4A, ATP4A* mean corr. ± STD 0.94±0.013], chief cells (CC) [*PGC, LIPF, AZGP1*, 0.89±0.013], gastric enteroendocrine cells (GEEC) [*ST18, INSM1, ARX*, 0.89±0.021], gastric mucous cells (GMC) [*LGALS4, VILL, CAPN8*, 0.94±0.008], mitotic cells (MTC) [*NCAPG, KIFC1, NCAPH*, 0.93±0.009], endothelial cells (EC) [*PECAM1, CDH5, ERG*, 0.89±0.013], fibroblasts (FB) [*PCOLCE, CLEC11A, MMP2*, 0.87±0.027], macrophages (MC) [*C1QB, FCGR3A, ITGB2*, 0.86±0.015], neutrophils (NP) [*CXCR2, FCGR3B, CXCR1*, 0.86±0.009], T-cells (TC) [*CD3E, CD2, CD3G*, 0.9±0.019] and plasma cells (PC) [*IGKC, JCHAIN, IGLC1*, 0.97±0.009]. (**ii**) A low corr. between *Ref*.*T*. across different cell type panels (Figure 1 C) (Table S1, Tab 1), indicating *specificity* (mean inter-panel corr. ± STD 0.08 ± 0.14) and (**iii**) A normal distribution of *Ref*.*T*. expression across the samples (Figure S1 A).

**Figure 1.**
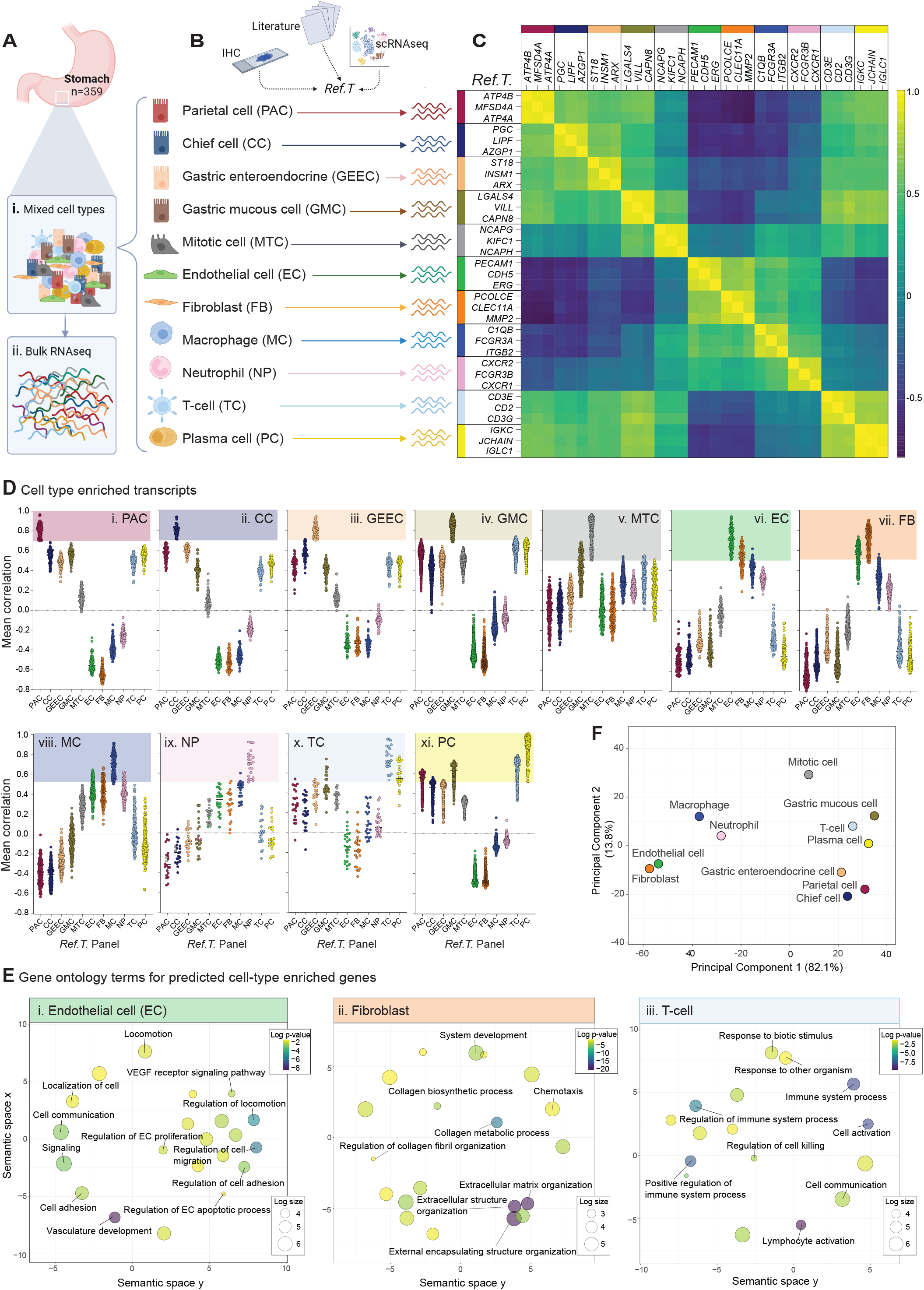
Integrative co-expression analysis can resolve constituent cell type identities from unfractionated human stomach tissue RNAseq data. (**A**) RNAseq data for 359 unfractionated human stomach samples were retrieved from GTEx V8. Each sample contained (i) mixed cell types, which contributed (ii) differing proportions of sequenced mRNA. (**B**) To profile cell type-enriched transcriptomes, constituent cell types were identified and candidate marker genes (’reference transcripts’ [*Ref.T*.]) for virtual tagging of each were selected, based on in house tissue protein profiling and/or existing literature and datasets. (**C**) Matrix of correlation coefficients between selected *Ref.T*. across the sample set. (**D**) Mean correlation coefficients of genes above designated thresholds for classification as cell-type enriched in stomach: (i) parietal cells [PC], (ii) chief cells [CC], (iii) gastric enteroendocrine cells [GEEC], (iv) gastric mucous cells [GMC], (v) mitotic cells [MTC], (vi) endothelial cells [EC], (vii) fibroblasts [FB], (viii) macrophages [MC], (ix) neutrophils [NP], (x) T-cells [TC], (xi) plasma cells [PC] with all *Ref.T*. panels. (**E**) Over-represented gene ontology terms among genes predicted to be: (i) endothelial cell, (ii) fibroblast or (iii) T-cell enriched. (**F**) Principal component analysis of correlation profiles of cell type enriched genes. See also Table S1 Tab 1 and 2.

#### Using reference transcript analysis to identify cell type-enriched genes

Correlation coefficients (corr.) between each selected *Ref*.*T*. and all other sequenced transcripts (>56,000) were calculated across stomach RNAseq samples. The proportion of cell types represented in each sample varies, due to biological and sampling variability, but ratios should remain consistent between constitutively expressed cell-enriched genes. Thus, a high corr. of a given transcript with all *Ref*.*T*. in only one cell type panel is consistent with enrichment in the corresponding cell type. For each cell type, a list of enriched genes was generated (Figure 1 D.i-xi) with inclusion based on: (**i**) the gene having the highest mean corr. with the *Ref*.*T*. panel representing a given cell type, over a specified threshold - indicated by a shaded box (Figure 1 D) and (**ii**) a differential between this value and the mean corr. with any other *Ref*.*T*. panel >0.15. This excluded genes that are co-enriched in two or more cell types, as we previously described (Norreen-Thorsen *et al*., 2022) (all data in Table S1, Tab 2). In some cases, genes had a high corr. value with more than one *Ref*.*T*. panel, e.g., those most highly correlating with the fibroblast *Ref*.*T*. panel (Figure 1 D.vii) showed elevated corr. with the *Ref*.*T*. panel for endothelial cells, and *vice versa* (Figure 1 D.vi). However, all cell type enriched genes were well separated when the individual minimum differential corr. were plotted against all other *Ref*.*T*. panels (Figure S1 B). Furthermore, gene ontology (GO) and reactome analysis (Ashburner *et al*., 2000; Gene Ontology, 2021) revealed that the over represented terms for these cell types were consistent with known functions e.g., for endothelial cells terms included ’*vascular development’* (FDR 3.7 ×10^−9^) and ’*angiogenesis’* (FDR 4.7 ×10^−9^) (Figure 1 E.i) for fibroblasts ’*extracellular structure organisation’* (FDR 3.6 ×10^−21^) and ’*collagen metabolic process’* (FDR 1.1 ×10^−13^) (Figure 1 E.ii) and for T-cells ’*T-cell activation*’ (FDR 1.3 ×10^−10^) and ’*lymphocyte activation*’ (FDR 2.0 ×10^−10^) (Figure 1 E.iii) (Table S1, Tab 8, 9 and 12). Principal component analysis of the corr. values of cell type-enriched genes revealed the largest variance was between stomach specific cell types vs. stromal/vasculature related ones (Figure 1 F).

### Stomach cell type enriched gene signatures

#### The majority of stomach cell type enriched genes are protein coding

1634 transcripts were predicted to be cell type-enriched (Figure 2 A and Table S1, Tab 2). Gastric mucous cells, plasma cells and mitotic cells had the highest number of predicted enriched genes (n=459, 275 and 179 respectively) (Figure 2 A.i, ii and iii). Of the other cell types found in all, or most, tissue types, fibroblasts and macrophages had the most enriched genes (n=203 and 161, respectively) (Figure 1 A.iv-v). Other stomach specialised cell types, parietal cells, chief cells and gastric enteroendocrine cells, had significantly fewer enriched genes (n=96, 66 and 56, respectively) (Figure 2 A.vi, viii and ix), and neutrophils and T-cells had the fewest overall (n=22 and 24, respectively) (Figure 2 A.x and xi). In all cases, the majority of cell type enriched transcripts were classified as protein coding (Yates *et al*., 2020), with the exception of plasma cells, in which immunoglobulin (IG) gene was the most common classification (Figure 2 A.ii). lncRNA were the most common type of non-coding cell type enriched transcript, with the exception of plasma cells, where immunoglobulin (IG) pseudogene was the most common non-coding classification (Figure 2 A.ii).

**Figure 2.**
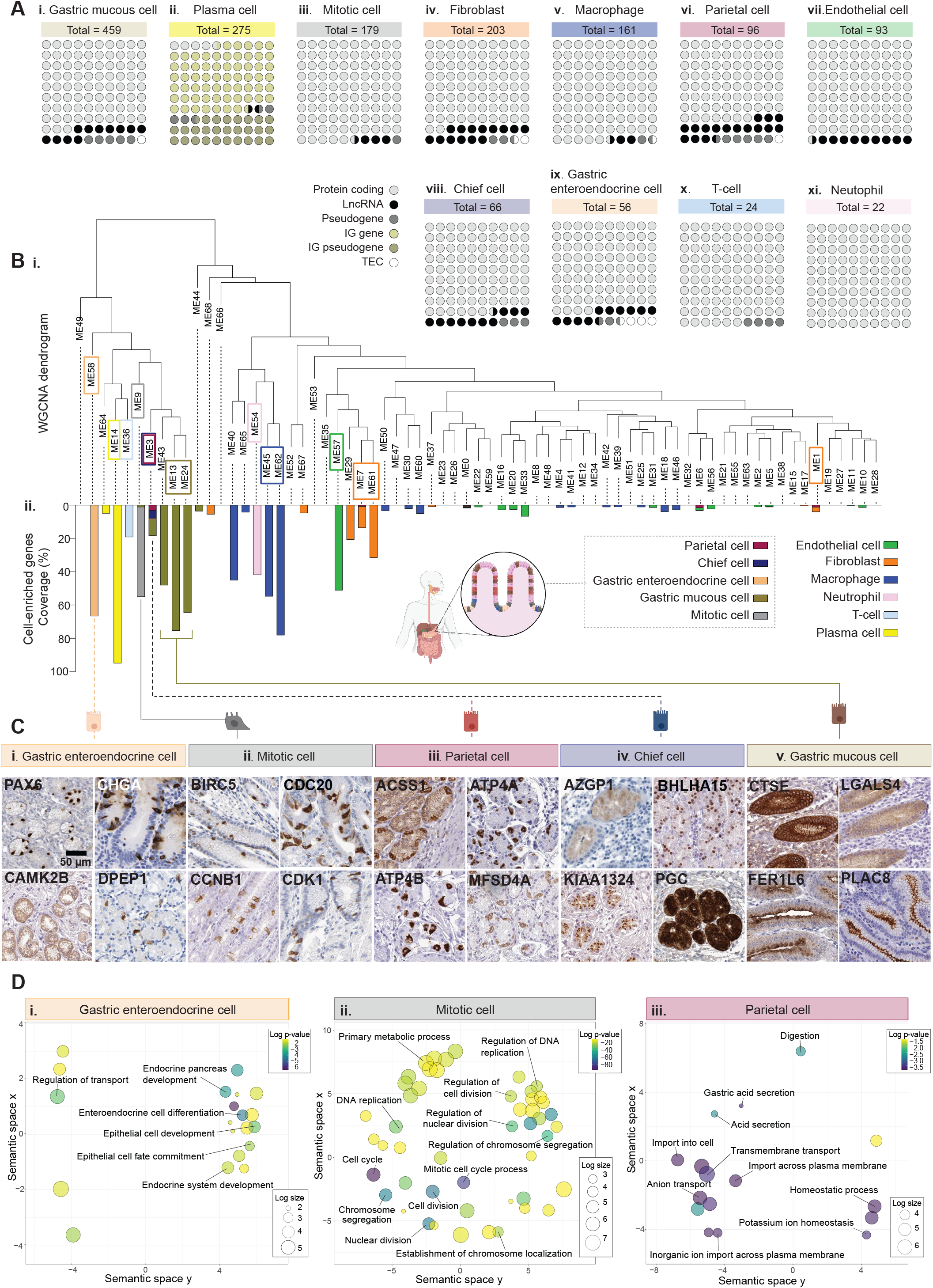
Integrative co-expression analysis of unfractionated RNAseq reveals enriched transcripts in human stomach cell types. (**A**) Total number and proportional representation of transcript class for cell type enriched genes in: (i) gastric mucous cells, (ii) plasma cells, (iii) mitotic cells, (iv) fibroblasts, (v) macrophages, (vi) parietal cells, (vii) endothelial cells, (viii) chief cells, (ix) gastric enteroendocrine cells, (x) T-cells and (xi) neutrophils. (**B**) RNAseq data for 359 unfractionated human stomach samples was subject to weighted correlation network analysis (WGCNA). (i) Coloured squares indicate cell type *Ref.T*. positions on resultant dendrogram. (ii) Coloured bars show distribution of protein coding transcripts classified as cell type-enriched across dendrogram groups. (**C**) Human stomach tissue profiling for proteins encoded by genes classified as: (i) gastric enteroendocrine cell, (ii) mitotic cell, (iii) parietal cell, (iv) chief cell or (v) gastric mucous cell enriched. (**D**) Over-represented gene ontology terms among genes predicted to be (i) gastric enteroendocrine cell, (ii) mitotic cell or (iii) parietal cell enriched. See also Table S1 Tab 2, 3, 5 and 6.

### Alternative analysis and protein profiling support cell-type classifications

#### Unsupervised weighted network correlation analysis is consistent with Ref.T. analysis

As our analysis is based on manually selected *Ref*.*T*. panels, cell-type classification is subject to an input bias. As a comparison, we subjected the same GTEx RNAseq dataset to a weighted network correlation analysis (WGCNA) (Langfelder and Horvath, 2008), an unbiased method that does not require any manual input or marker gene selection. WGNCA generates corr. coefficients between all transcripts and subsequently clusters them into related groups, based on expression similarity (Figure 2 B). In general, *Ref*.*T*. belonging to the same cell type panel were found in the same WCGNA cluster (Figure 2 B.i, coloured boxes represent *Ref*.*T*. locations), e.g., gastric enteroendocrine cells (cluster 58) or adjacent clusters on the same branch, e.g., gastric mucous cells (clusters 13 and 24) and macrophages (clusters 45 and 62) (Figure 2 B.i). Protein coding genes that we predicted to be cell type enriched were predominantly clustered into the same WGCNA group as the corresponding *Ref*.*T*., or into adjacent groups on the same branch, consistent with our classifications (Figure 2 B.ii). Genes in the *Ref*.*T*. panels representing parietal and chief cells appeared in the same large group (cluster 3) (Figure 2 B.ii), as were the genes in the respective predicted enriched gene lists, despite clear separation in our *Ref*.*T* based method (Figure 1 C, D). Despite the lack of separation for the enriched gene signatures for parietal and chief cells by WGNCA, each contained several well described marker genes for the respective cell type, e.g., *GIF, SLC26A7* (parietal) and *PGA4, SLC1A2* (chief cell). Indeed, we have previously shown that *Ref*.*T*. based analysis can have a higher sensitivity than WGNCA for cell type gene enrichment analysis (Dusart *et al*., 2019). Stomach tissue protein profiling revealed staining consistent with expression in the respective cell types for proteins encoded by genes predicted to be gastric enteroendocrine cell (Figure 2 C.i), mitotic cell (Figure 2 C.ii), parietal cell (Figure 2 C.iii), chief cell (Figure 2 C.iv) or gastric mucous cell (Figure 2 C.v) enriched. GO and reactome analysis (Ashburner *et al*., 2000; Gene Ontology, 2021) revealed over represented terms for predicted stomach specialised cell type enriched genes were consistent with known cell functions e.g., for gastric enteroendocrine cells *’enteroendocrine cell differentiation’* (GO; FDR 4.06 × 10^−5^) and *’regulation of gene expression in endocrine-committed (NEUROG3+) progenitor cells’* (reactome; FDR 5.45 × 10^−4^) (Figure 2 D.i), for mitotic cells ’*mitotic cell cycle*’ (GO; FDR 1.03×10^−74^) and ’*cell cycle*’ (reactome; FDR 5.05×10^−69^) (Figure 2 D.ii) and for parietal cells ‘*gastric acid secretion*’ (GO; FDR 2.25 × 10^−4^) and ’*transport of small molecules*’ (reactome; FDR 1.78 × 10^−4^) (Figure 2 D.iii), (for all cell types see Table S1, Tab 3-13).

#### Stomach cell type gene enrichment signatures

Figure 3 shows up to the top 50 most enriched protein coding enriched transcripts for each cell type, ranked by highest corr. with the relevant *Ref*.*T*. panel (Figure 3 A.i-K.i), with differential corr. values and expression levels in the bulk RNAseq dataset (mean TPM). Mean TPM levels were generally highest for genes predicted to be enriched in parietal cells (Figure 3 A.i), chief cells (Figure 3 B.i), gastric mucous cells (Figure 3 D.i), fibroblasts (Figure 3 G.i) and plasma cells (Figure 3 K.i), and lowest for those in mitotic cells (Figure 3 E.i), neutrophils (Figure 3 I.i) and T-cells (Figure 3 J.i). This likely reflects differing numbers of each given cell type with the samples, however, as a range of expression values are observed within each given cell type, there is likely also individual gene variation in factors such as regulation and transcript stability. The highest differential values, and thus relative uniqueness among the profiled cell types, was observed for mitotic cell enriched genes (Figure 3 E.i), most of which have well studied roles in the regulation of the cell cycle, such as *TOP2A* and *BUB1B*. For all other cell types, top enriched genes included both known cell type specific genes, together with those that have not been previously reported as such, e.g., *PECAM1* and *SHE* were both predicted to be endothelial cell enriched (Figure 3 F.i); *PECAM1* is a commonly used marker gene for this cell type, whilst there are no existing reports for the selective expression of *SHE* in this context. Tissue profiling for proteins encoded by representative cell type enriched genes showed expression consistent with our classifications (Figure 3 A.ii-K.ii).

**Figure 3.**
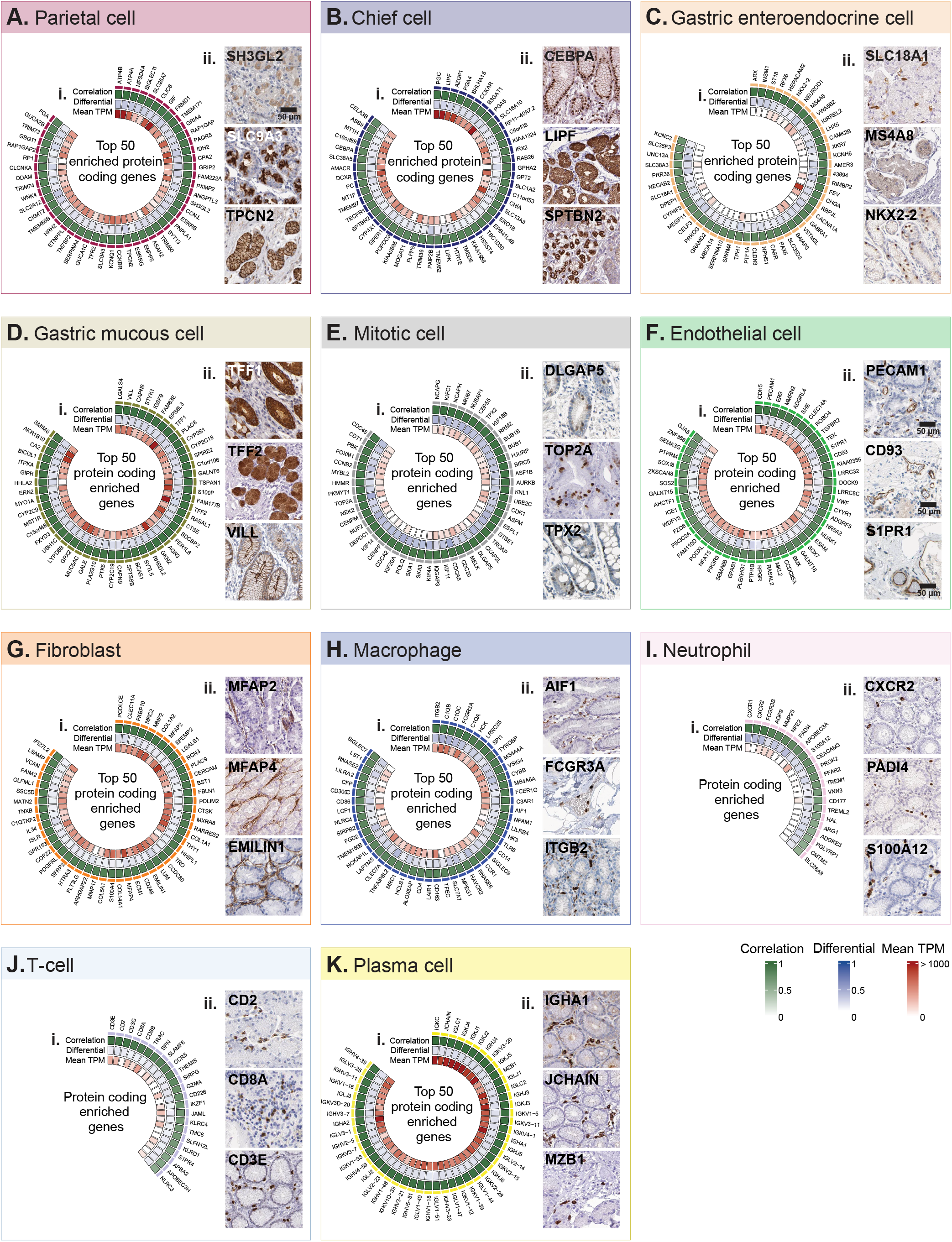
Protein coding gene signatures of human stomach cell types. Cell type-enriched protein coding transcripts in: (**A**) parietal cells, (**B**) chief cells, (**C**) gastric enteroendocrine cells, (**D**) gastric mucous cells, (**E**) mitotic cells, (**F**) endothelial cells, (**G**) fibroblasts, (**H**) macrophages, (**I**) neutrophils (**J**) T-cells and (**K**) plasma cells, showing: (i) correlation coefficient with the cell type *Ref.T*. panel, differential correlation scores (correlation with cell type *Ref.T*., panel minus max correlation with any other *Ref.T*. panel) and mean expression in bulk RNAseq. (ii) Human stomach tissue protein profiling for selected cell type enriched genes. See also Table S1 Tab 2.

#### Ref.T. analysis can predict source of stomach enriched protein coding genes

Genes with enriched expression in the human stomach vs. other tissue types can be identified by a comparative analysis of unfractionated tissue RNAseq data. We extracted the top 200 human stomach-enriched genes from the Human Protein Atlas (HPA) (Uhlen *et al*., 2015) and GTEx project (Consortium, 2015), through the Harminozome database (Rouillard *et al*., 2016) (Figure 4). Of the 78 genes classified as stomach-enriched in both datasets, 46/78 (59.0%) were classified as cell type enriched in our analysis; 28/46 (61.0%) in gastric mucous cells, 11/46 (24.0%) in parietal cells, 6/46 (13.0%) in chief cells, and 1/46 (2.2%) in gastric enteroendocrine cells (Figure 4 B.i and B.ii, respectively, large symbols). Of those not classified as cell type-enriched in our analysis (n=32), 11/32 (34.4%), only narrowly failed to reach one of the thresholds for classification as either parietal-, chief- or gastric mucous cell-enriched (Figure 4 B.i and B.ii, medium symbols). The majority of the remaining genes most highly correlated with *Ref.T*. panel representing one, or more, of the same cell types; parietal, chief or gastric mucous, but were excluded from the cell-type classifications due to shared enrichment. None of the stomach-enriched genes were predicted to be enriched in any cell type found across multiple tissue types, such as endothelial or immune cells, consistent with the lack of specificity of these cell type to the stomach. Thus, our analysis indicates that most stomach-tissue enriched genes are primarily expressed in gastric mucous, parietal or chief cells.

**Figure 4.**
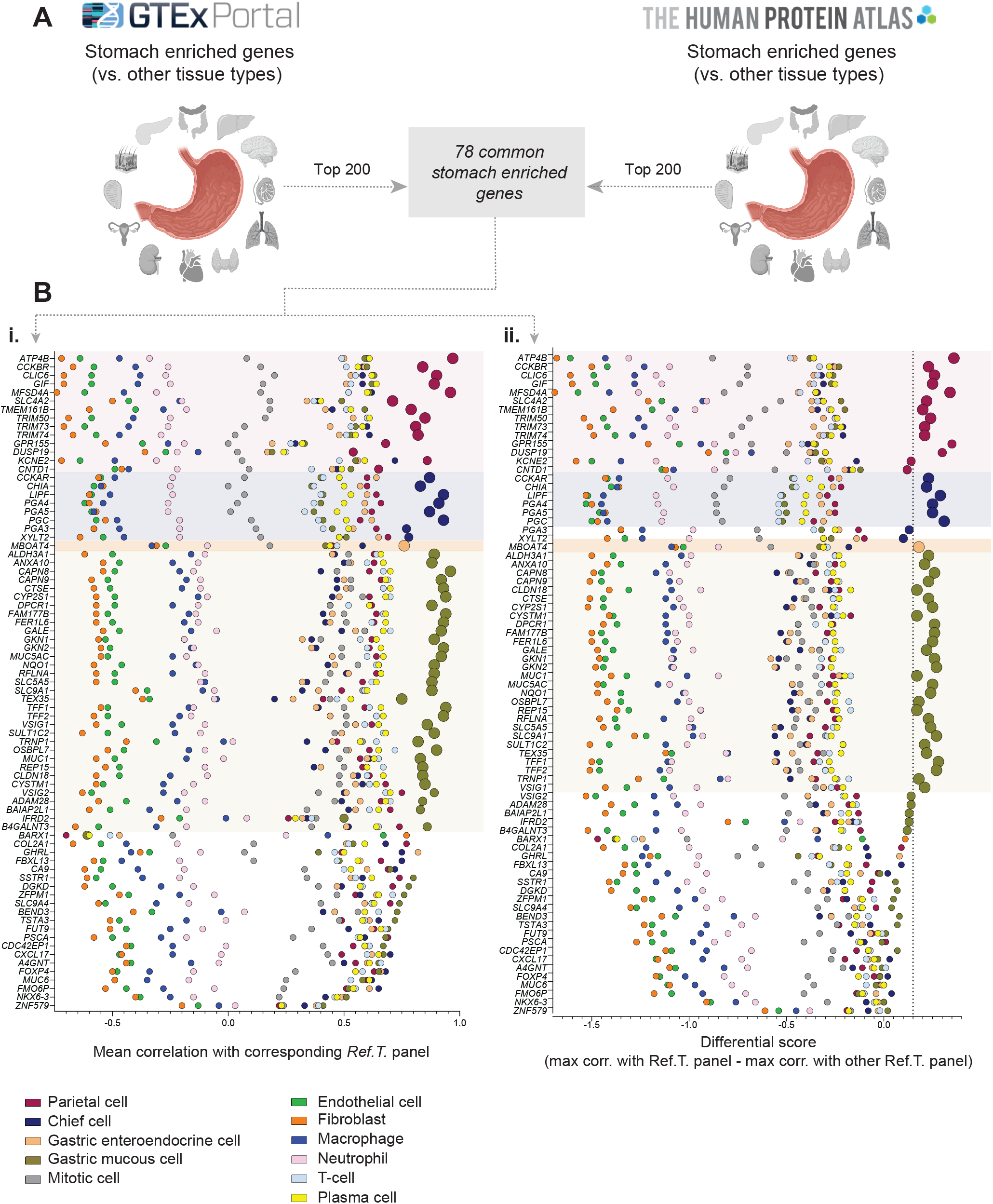
Gastric mucous cells, parietal cells and chief cells are the primary source of stomach tissue enriched genes. (**A**) The top 200 stomach enriched genes (vs. other tissue types) in RNAseq data from the GTEx Portal or Human Protein Atlas (HPA) were compared to identify genes common to both datasets (n=78). For each, the following was plotted: (**B**) (i) the mean correlation with each cell type *Ref.T*. panel, and (ii) the differential value vs. the next most highly correlating *Ref.T*. panel (dotted line indicates threshold for classification as cell type enriched). Enlarged circles represent genes with predicted cell type enrichment.

### Cell-type enriched non-coding transcripts in stomach

A total of 272 non-coding transcripts were identified as cell-type enriched in the stomach (Figure 5 A), the greatest number of which were in plasma cells, gastric mucous cells or fibroblasts (n=89, 79 and 37, respectively). When the sample set was analysed by WGNCA (Figure 5 B.i) non-coding transcripts that we predicted to be cell type enriched predominantly clustered into the same WGCNA group as the corresponding *Ref.T*., or into adjacent groups on the same branch (Figure 5 B.ii). Up to the top 50 non-coding enriched genes in gastric enteroendocrine cells (Figure 5 C.i), gastric mucous cells (Figure 5 D.i), fibroblasts (Figure 5 E.i), chief cells (Figure 6 A.i), parietal cells (Figure 6 B.i), plasma cells (Figure 6 C.i), and endothelial cells (Figure 6 D.i), ranked by corr. with the relevant Ref.T panel, are displayed with differential corr. values vs. other profiled cell types, expression in the bulk RNAseq data (mean TMP) and transcript type. In all cell types, with the exception of plasma cells, where the most common type of enriched non-coding gene was IG pseudogene (Figure 6 C.i), long non-coding RNAs made up the majority of the predicted enriched transcripts. Generally, gastric mucous cell (Figure 5 D.i) and fibroblast (Figure 5 E.i) enriched non-coding genes were expressed at the highest levels in the stomach bulk RNAseq. This likely reflects the differing numbers of each given cell type within the samples, but the intra-cell type variation also indicates individual gene regulation.

**Figure 5.**
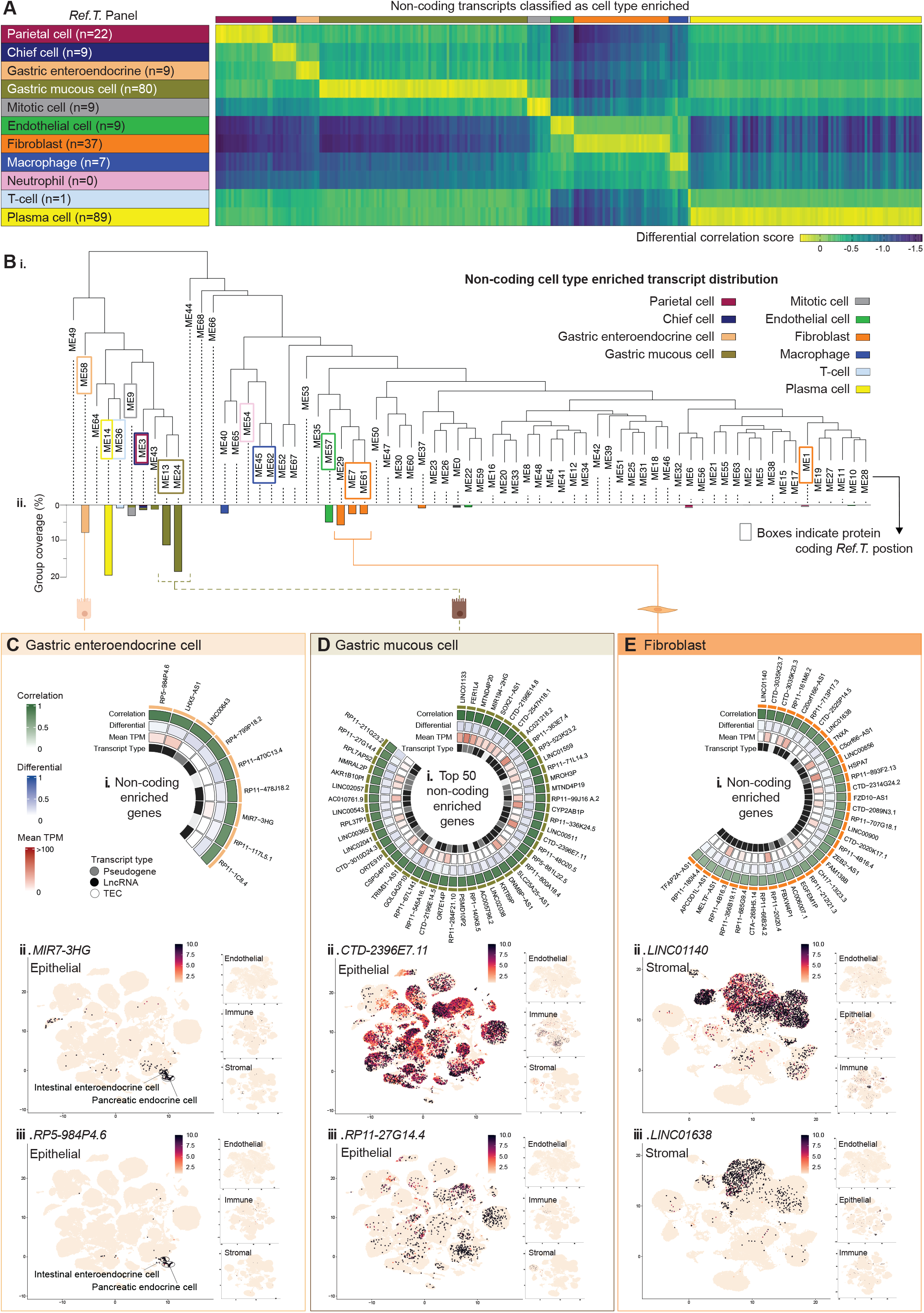
Non-coding gene signatures of human stomach cell types. **(A)** Heat map of non-coding transcripts predicted to be cell type enriched, showing differential score between mean correlation coefficient with the corresponding *Ref.T*. panel vs. highest mean correlation coefficient amongst the other *Ref.T*. panels. (**B**) RNAseq data for 359 unfractionated human stomach samples was subject to weighted correlation network analysis (WGCNA). (i) Coloured squares indicate cell type *Ref.T*. positions on resultant dendrogram. (ii) Coloured bars show distribution of non-coding genes classified as cell type-enriched across dendrogram groups. Non-coding gene enrichment signatures for: (**C**) gastric enteroendocrine cells, (**D**) gastric mucous cells and (**E**) fibroblasts, detailing: (i) up to the top 50 cell type enriched non-coding genes, showing correlation coefficients with the *Ref.T*. panel, differential scores (correlation with corresponding cell type *Ref.T*., panel minus max correlation with any other *Ref.T*. panel), mean expression in bulk RNAseq and transcript type. (ii and iii) scRNAseq data from analysis of epithelial, endothelial, immune or stromal cell compartments across 24 human tissues was sourced from Tabula Sapiens (Tabula Sapiens et al., 2022), and used to generate UMAP plots showing the expression profiles of example cell type enriched non-coding genes. The largest plot shows the compartment with the highest expression. See also Table S1 Tab 2 and Figure S2 (for all UMAP plot annotations).

**Figure 6.**
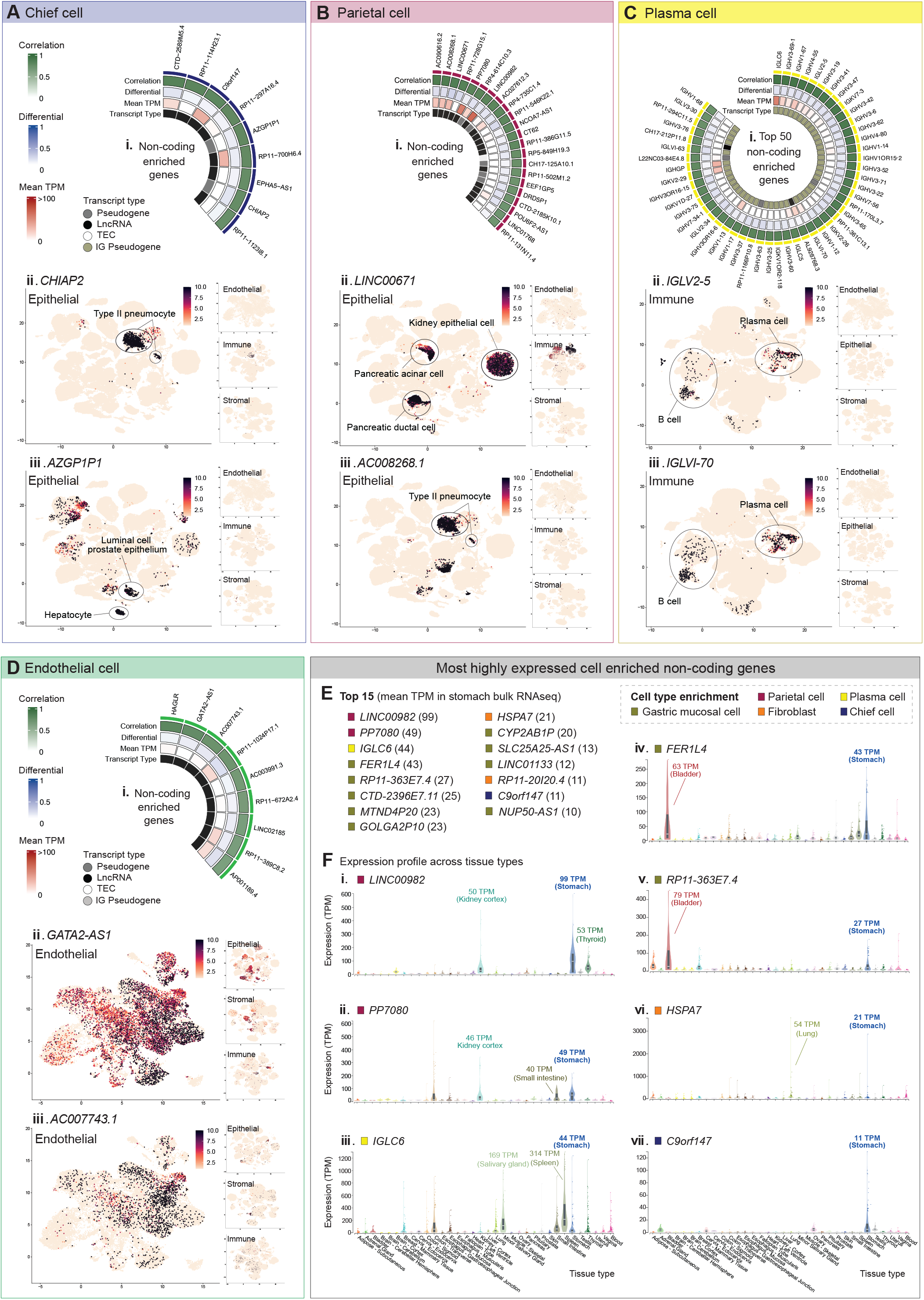
Core non-coding gene signatures of human stomach cell types and tissue distribution patterns. Non-coding gene enrichment signatures for: (**A**) chief cells, (**B**) parietal cells, (**C**) plasma cells and (**D**) endothelial cells, detailing (i) up to the top 50 cell type enriched non-coding genes, showing correlation coefficients with the *Ref.T*. panel, differential scores (correlation with corresponding cell type *Ref.T*., panel minus max correlation with any other *Ref.T*. panel), mean expression in bulk RNAseq and transcript type. (ii and iii) scRNAseq data from analysis of epithelial, endothelial, immune, or stromal cell compartments across 24 human tissues was sourced from Tabula Sapiens (Tabula Sapiens et al., 2022), and used to generate UMAP plots showing the expression profiles of example cell type enriched non-coding genes. The largest plot shows the compartment with the highest expression. (**E**) The top 50 most highly expressed cell type enriched non-coding genes in stomach bulk RNAseq. (**F**) Expression of genes classified as enriched in parietal cells: (i) *LINC00982* and (ii) *PP7080*, plasma cells: (iii) *IGLC6*, gastric mucous cells: (vi) *FER1L4* and (v) *RP11-363E7.4*, fibroblasts: (vi) *HSPA7* and chief cells: (vii) *C9orf147*, in bulk RNAseq of different human organs. Mean TMP expression is annotated for selected organs on each plot. See also Table S1 Tab 2 and Figure S2 (for all UMAP plot annotations).

There is currently no existing dataset of non-coding enriched genes in stomach cell types that could be used to validate our predictions. However, we sourced scRNAseq data from the analysis of 24 tissue types in Tabula sapiens (Tabula Sapiens *et al*., 2022) (data for stomach was not available) that had been classified into endothelial, epithelial, immune and stromal cell functional compartments (for Tabula sapiens UMAP cell type classifications see Figure S2). We generated UMAP plots for each of these compartments to determine expression profiles for selected non-coding genes that we predicted to be cell type enriched. The predicted gastric enteroendocrine enriched genes *MIR7-3HG* and *RP5-984P4.6* were expressed only in the epithelial cell compartment, specifically in the clusters annotated as intestinal enteroendocrine and pancreatic alpha and beta cells (Figure 5 C.ii and iii), consistent with a specialised role in endocrine cells, not only in the stomach, but also in the pancreas and other parts of the GI tract. The predicted gastric mucous cell enriched genes *CTD-2396E7.11* and *RP11-27G14.4* were widely expressed in the epithelial compartment, but not in the endothelial, immune, or stromal cell compartments (Figure 5 D.ii and iii). The predicted fibroblast enriched genes *LIN01140* and *LINC01638* were expressed predominantly in the stromal cell compartment (Figure 5 E.ii and iii), also consistent with our classifications. Genes predicted to be chief cell enriched, *CHIAP2* and *AZGP1P1* (Figure 6 A.ii and iii), and parietal cell enriched, *LINC00671* and *AC008268.1* (Figure 6 B.ii and iii), were predominantly expressed in the epithelial compartment. The type of epithelial cell in which the genes were expressed varied, e.g., the chief cell enriched gene *CHIAP2* (Figure 6 A.ii) was only expressed in type II pneumocytes from the lung; one could speculate that this indicates a shared specialised function of these specific cell types, as both are highly specialised secretory cells for their respective tissues. The predicted plasma cell enriched genes *IGLV2-5* and *IGLVI-70* were expressed only in the immune cell compartment (Figure 6 C.ii and iii) in clusters annotated as either plasma cells or B-cells, and those predicted to be endothelial cell enriched, *GATA-2-AS1* and *AC007743.1* (Figure 6 D.ii and iii) were predominantly expressed in the endothelial compartment. Thus, the Tabula sapiens scRNAseq data provides supportive evidence for our cell type classifications, despite the lack of stomach cell type analysis in this dataset.

Of those non-coding genes that we classified as cell type enriched, 15 had relatively high expression in the bulk RNAseq stomach samples (mean TPM >10), the majority of which were predicted to be gastric mucous cell enriched (Figure 6 E). To determine the expression profile of these genes in different organ types, we sourced data from bulk RNAseq of other tissues in GTEx (Figure 6 F). The most highly expressed parietal cell enriched non-coding transcripts, *LINC00982* and *PP7080* (mean TPM 99 and 49, respectively) both had high relative expression in stomach tissue (Figure 6 F.i and ii), consistent with a specialised function in this organ. *IGLC6*, the most highly expressed non-coding transcript we predicted to be enriched in plasma cells was highly expressed in spleen and salivary gland; tissues that contain high numbers of plasma cells (Figure 6 F.iii). The most highly expressed non-coding transcripts we predicted to be enriched in gastric mucous cells, *FER1L4* and *RP11-363E7.4*, both had high relative expression in stomach and bladder (Figure 6 F.iv and v); one could speculate these genes have specific functions in the mucous cells found in these tissue types. *HSPA7*, the most highly expressed predicted fibroblast enriched gene had variable expression across tissue types (Figure 6 F.vi), consistent with the ubiquitous presence of this cell type across organs, whilst the most highly chief cell enriched transcript, *C9orf147*, had high relative expression only in stomach tissue (Figure 6 F.vii). Thus, the most highly expressed non-coding transcripts predicted to be enriched in the stomach specialised cell types were predominantly detected in stomach tissue, consistent with a specialised function here. Conversely, those predicted to be enriched in less specialised cell types were more broadly expressed across tissue types, consistent with a common cell type function. All data for non-coding transcripts can be searched via the web portal https://cell-enrichment.shinyapps.io/noncoding_stomach/.

#### Comparison of predicted sex-specific stomach cell type enriched genes

We performed a subset analysis of the stomach RNAseq dataset (male n=227, female n=132,), to identify sex-specific cell type enriched genes. Similar to the full dataset, intra-panel cell type *Ref.T*. correlated well in single-sex sample subsets (all >0.84) (Table S2, Tab 1, Table A and B). Cell type enriched genes were calculated as for the whole dataset. To compare gene enrichment profiles in males and females, the following was calculated for any gene that was classified as cell type enriched in either subset: (**i**) the ‘*differential correlation score*’, defined as the difference between the mean corr. coefficient with the cell type *Ref.T*, in the male and female sample subsets, (**ii**) the ‘*enrichment ranking*’, based on the mean corr. value with the *Ref.T*. panel (rank 1 = highest corr.). Cell profiles were mainly comparable between sexes, for both stomach specialised cell types (Figure 7 Ai-iv) and others (Figure S3 A-G) (transcripts enriched in *both* males and females represented by square symbols). For those genes classified as enriched *only* in males or females (represented by differently coloured triangle and circle symbols, respectively), most had differential corr. scores close to zero; indicating that they fell marginally below the designated threshold for classification as enriched in the other sex. A small number of distinct male-only enriched transcripts were identified in chief cells; *ARSFP1, TBL1Y* and *RP11-115H13.1* (Figure 7 A.iv), all of which were Y-linked, with expression levels above background level only in male samples (Figure 7 Bi-iii). As described above, we sourced scRNAseq data from Tabula sapiens (Tabula Sapiens *et al*., 2022) for cells classified as endothelial, epithelial, immune or stromal (Figure S2). We generated UMAP plots (using cell data from male donors only) to show expression profiles of the male-only chief cell enriched genes. *ARSFP1* was detected only at low levels in the epithelial compartment (Figure 7 C.i), whilst *TBL1Y* (Figure 7 C.ii) and *RP11-115H13.1* (Figure 7 C.iii) had strikingly similar expression profiles, with the highest levels in both cases detected in prostate epithelial cells. All 3 male-only chief cell enrichened genes had low/no expression in the endothelial, immune or stromal compartments (Figure 7 Ci-iii). To determine the broad expression profile of the most highly expressed non-coding enriched genes across organs (from male donors), we sourced data from GTEx (Figure 7 D). *ARSFP1* had enhanced expression only in the stomach and esophagus (Figure 7 D.i); both of which are tissue types not included in the Tabula sapiens dataset, consistent with the low detection observed there. *TBL1Y* and *RP11-115H13.1* had similar expression profiles across tissue types, with enhanced expression in thyroid (which was also absent from the Tabula Sapiens dataset) followed by prostate; in keeping with the high expression observed in prostate epithelial cells in the scRNAseq (Figure 7 D.ii-iii). Thus, one could speculate that male-only chief cell enriched gene *ARSFP1* has a stomach specific function, whilst *TBL1Y* and *RP11-115H13.1* appear to be co-expressed also in cell types outside the stomach, suggesting a broader function in multiple cell types.

**Figure 7.**
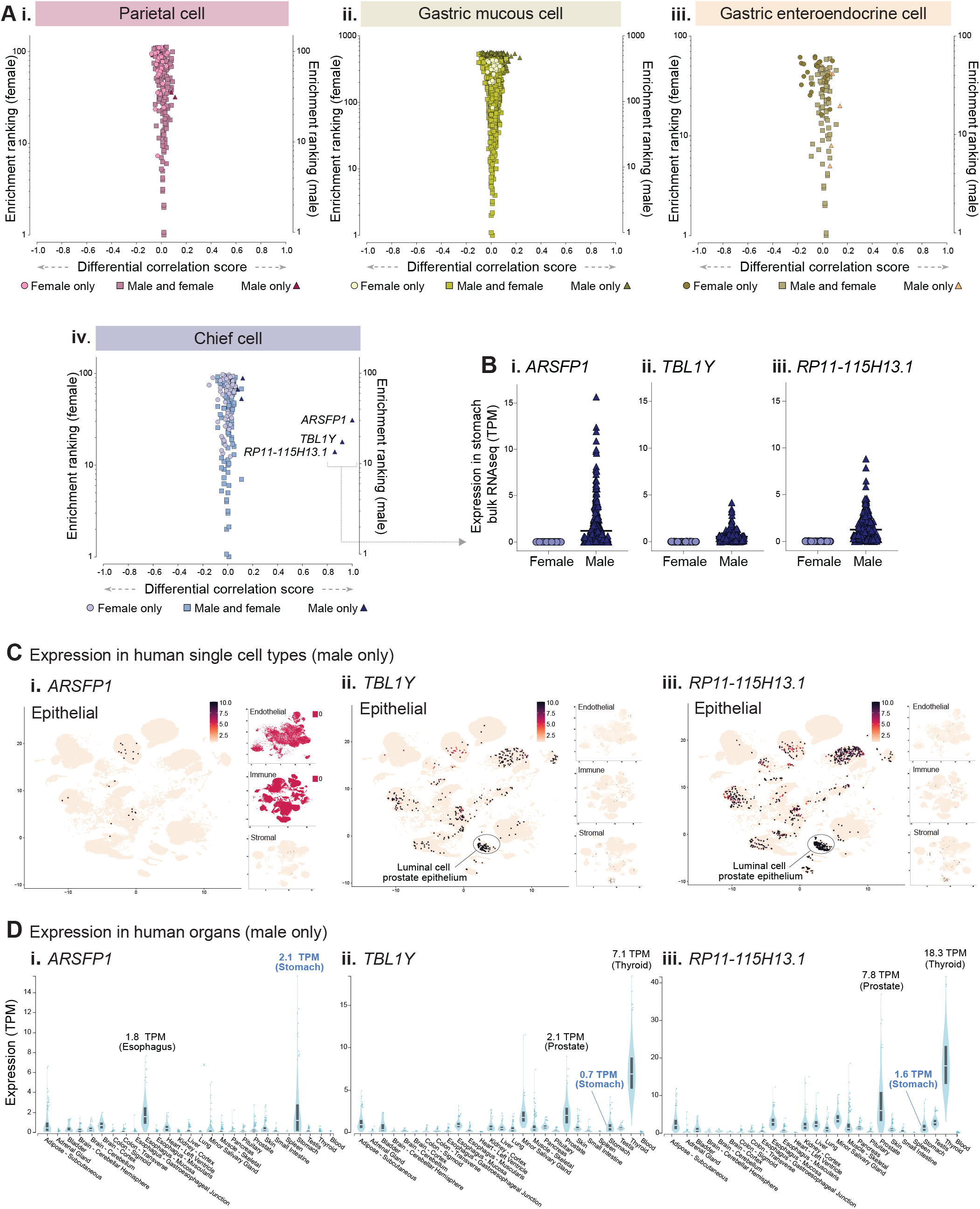
Identification of sex-specific cell-enriched transcripts in human stomach tissue. (**A**) Human stomach RNAseq data (n=359 individuals) was retrieved from GTEx V8 and divided into female (n=132) and male (n=227) subgroups before classification of cell type-enriched transcripts. For transcripts classified as: (i) parietal, (ii) gastric mucous, (iii) gastric enteroendocrine or (vi) chief cell enriched transcripts. In either sex, the ’*sex differential corr. score*’ (difference between mean corr. with the *Ref.T*. panel in females vs. males) was plotted vs. ‘enrichment ranking’ (position in each respective enriched list, highest corr. = rank 1). On each plot, transcripts enriched in *both* females and males are represented by common-coloured square symbols, and transcripts classified as enriched *only* in females or males are represented by differently coloured circle and triangle symbols, respectively. (**B**) Expression in female or male samples for transcripts classified as male-only enriched in chief cells: (i) *ARSFP1*, (iii) *TBL1Y* and (iii) *RP11-115H13.1*. (**C**) scRNAseq data from analysis of epithelial, endothelial, immune or stromal cell compartments across human tissues from male donors was sourced from Tabula Sapiens (Tabula Sapiens et al., 2022), and used to generate UMAP plots showing the expression profiles of: (i) *ARSFP1*, (iii) *TBL1Y* and (iii) *RP11-115H13.1*. (**D**) Expression of: (i) *ARSFP1*, (iii) *TBL1Y* and (iii) *RP11-115H13.1* in bulk RNAseq of different human organs from male donors. The largest plot shows the compartment with the highest expression. Mean expression is annotated for selected organs on each plot. See also Table S2 Tab 1, Figure S2 (for all UMAP plot annotations) and Figure S3.

## DISCUSSION

Here, we present a genome wide cell type enriched transcriptome atlas for the human stomach, using our previously described method to resolve unfractionated tissue RNAseq data to the cell type level (Butler *et al*., 2016; Dusart *et al*., 2019; Norreen-Thorsen *et al*., 2022). Our method circumvents some challenges associated with scRNAseq analysis, including issues associated with cell isolation, material amplification (Shapiro, Biezuner and Linnarsson, 2013; Grün and van Oudenaarden, 2015; Gawad, Koh and Quake, 2016) and induction of expression artefacts, due to loss of tissue specific cues or processing (O’Flanagan *et al*., 2019). Our analysis incorporates a high number of biological replicates, reducing the impact of individual variation and allowing for well powered subgroup comparisons e.g., female vs. male. As data for gene enrichment signatures of stomach cell types are lacking in the existing literature, with this organ absent from large scale single cell sequencing (scRNAseq) initiatives, such as Tabula Sapiens (Tabula Sapiens *et al*., 2022) and the Human Cell Atlas (Regev *et al*., 2017) our study provides a useful resource, which can be searched on a gene-by-gene basis on the human protein atlas (www.proteinatlas.org/humanproteome/tissue+cell+type/stomach) or https://cell-enrichment.shinyapps.io/noncoding_stomach/, for protein coding and non-coding genes, respectively.

Of the 11 cell types we profiled in the stomach, gastric mucous cells had the highest number of predicted enriched genes, which included those encoding for proteins with known cell type specific functions, such as in mucosal defence, e.g., *CAPN8, CAPN9* (Hata *et al*., 2010), *GKN1* (Choi *et al*., 2013), *MUC13* (Ja *et al*., 2020), *TFF1* and *TFF2* (Aihara, Engevik and Montrose, 2017) and lipid metabolism, e.g., *PLPP2* (Hooks, Ragan and Lynch, 1998), *PPARG* (Kang *et al*., 2015) and *PLA2G10* (Hanasaki *et al*., 2002). In addition, several genes we identified have no reported role in this cell type, including *FAM83E, CYP2S1* and *PLAC8*. Predicted gastric enteroendocrine enriched genes also included those with known cell type function, such as *PAX6*, which controls endocrine cell differentiation (Beucher *et al*., 2012), and proglucagon (Hill, Asa and Drucker, 1999) and gastric inhibitory polypeptide (Fujita *et al*., 2008) production, *CAMK2B*, which is involved in intracellular calcium signalling (Tsakmaki *et al*., 2020), and the neuroendocrine secretory protein *CHGA* (Goldspink, Reimann and Gribble, 2018). Other predicted gastric enteroendocrine enriched genes had not been described in gastric enteroendocrine cells previously, such as *LHX5, SERPINA10* and *KCNH6. LHX5* has mainly been studied in the context of neuronal development (Zhao *et al*., 1999; Pillai *et al*., 2007), but in the GTEx database the only tissue type, outside the brain, where *LHX5* had elevated expression compared to others was the stomach (Consortium, 2015), thus, one could speculate that this gene also has a specific functional role here. *SERPINA10* was previously identified as a biomarker for gastrointestinal neuroendocrine carcinoma (Leja *et al*., 2009), and *KCNH6* has a role in the regulation of insulin secretion in the pancreas (J.-K. Yang *et al*., 2018); both consistent with our prediction that these genes have an endocrine cell enriched profile.

Many genes we predicted to be parietal cell enriched were well known markers of this cell type, such as *GIF* (Alpers and Russell-Jones, 2013) and *SLC26A7* (Petrovic *et al*., 2003). However, others had no reported cell type specific expression or function, such as *ACSS1*, a mitochondrial matrix protein functioning as a catalyst of acetyl-CoA synthesis (Schwer *et al*., 2006) and *MFSD4*, a marker for hepatic metastasis in gastric cancer (Shimizu, Kanda and Kodera, 2018). Our classifications were supported by a scRNAseq study that showed elevated expression of *ACSS1* and *MFSD4* in parietal cells vs. other stomach epithelial cells (Busslinger *et al*., 2021). Other predicted enriched genes for which a function in parietal cells has not yet been described included *SLC12A3, ETNPPL, FNDC10, TUBA3C, TRIM73, TRIM74* and *CLCNKA*. Chief cell enriched genes included *BHLHA15*, a known chief cell marker (Lennerz *et al*., 2010) and *KIAA1324*, which is required for chief cell secretory granule maturation (Cho, Park and Mills, 2022). Novel predicted chief cell enriched genes included the orphan receptor *GPR150*, a G-protein coupled receptor in which aberrant methylation has been linked to ovarian cancer (Cai *et al*., 2007), *MOGAT1*, a monoacylglycerol acyltransferase that functions in the absorption of dietary fat in the intestine (Yen *et al*., 2002) and *LIPK*, previously identified in the epidermis with a function in lipid metabolism (Toulza *et al*., 2007).

Whilst there is no existing database of non-coding gene enrichment profiles in the cell types of the stomach, and a lack of information regarding the function of any such genes in normal tissue, increasing evidence of the involvement of non-coding transcripts in the development of gastric cancer (Li *et al*., 2014; Gao *et al*., 2020; Ghafouri-Fard and Taheri, 2020) and associated drug resistance (Wei *et al*., 2020) indicates that this transcript class has important functions in this tissue type. Of the stomach specialised cell types we profiled, gastric mucous cells had the highest number of predicted enriched non-coding genes, which included several antisense transcripts to corresponding gastric mucous cell enriched protein coding genes, such as *FER1L6-AS1, SOX21-AS1* and *TRIM31-AS1*, suggesting a local regulation of gene transcription. Many gastric mucous cell enriched non-coding genes were expressed at relatively high levels, compared to other non-coding transcripts in the same or other cell types, including *LINC01133, FER1L4, RP11-363E7.4* and *CTD-2396E7.11. LINC01133* and the pseudogene *FER1L4* are inhibitors of gastric cancer progression, with reduced expression associated with a more aggressive tumour phenotype (Xia *et al*., 2015; X.-Z. Yang *et al*., 2018). To date, there is a single publication on *RP11-363E7.4*, where a genome wide screen of gastric cancer samples identified it as a key regulator of disease progression, with higher expression associated with overall survival (Wang *et al*., 2018). All the aforementioned studies were based on analysis of bulk RNAseq cancer samples, and the cell type in which these genes primarily function in healthy tissue is not reported; our data strongly indicates that this site is the mucous cell compartment. *CTD-2396E7.11* has not been described in the context of gastric cancer, but it was identified as one of four hub lncRNAs associated with reduced colon adenocarcinoma progression (Jiang, Tan and Zhang, 2019). As this tumour type also arises from the mucosa, on could speculate *CTD-2396E7.11* has a similar expression profile in healthy colon tissue. *LIN00982*, the highest expressed of all classified genes, was enriched in parietal cells and had, similar to those discussed above been shown to have a role in the inhibition of gastric cancer progression (Zheng *et al*., 2021).

Examples of non-coding genes we predicted to have gastric enteroendocrine cell enriched expression included *MIR7-3HG, RP5-984P4.6* and *RP11-470C13.4*. The selective expression of these genes in pancreatic and intestinal endocrine cells (Tabula Sapiens *et al*., 2022), is consistent with them having a conserved endocrine function. *MIR7-3HG* can act as an autophagy inhibitor (Capizzi *et al*., 2017), but there are no reports of its function in an endocrine context. *RP5-984P4.6* and *RP11-470C13.4* are currently completely uncharacterised. Other gastric enteroendocrine cell enriched non-coding genes included *LHX5-AS1*, an antisense transcript to the gastric enteroendocrine cell enriched corresponding protein coding gene.

Despite reported differences in stomach function between males and females, such as in speed of gastric emptying (Datz, Christian and Moore, 1987), gastrointestinal motility (Al-Shboul, 2016), incidence of gastric cancer (Lou *et al*., 2020) and in gastric cancer survival (Li *et al*., 2020), there are no studies of sex differences between stomach cell-type gene enrichment profiles. We found that global cell type gene enrichment signatures were similar between sexes, but we did identify 3 male-only chief cell enriched genes - *ARSFP1, RP11-115H13.1* and *TBL1Y*, all of which were Y-linked (Kirsch *et al*., 2004; Yan *et al*., 2005). In the GTEx database, the pseudogene *ARSFP1* was most highly expressed in male stomach samples, compared to the other 53 tissue types profiled from males (Consortium, 2015), supportive of a currently unknown sex and tissue specific role, and consistent with our predicted enrichment in a stomach-specific cell type in males. Although it is often assumed that pseudogenes lack function, recent studies have shown that they can have key roles, functioning as antisense, interference or competing endogenous transcripts (Pink *et al*., 2011; Kovalenko and Patrushev, 2018; Cheetham, Faulkner and Dinger, 2020). *RP11-115H13.1* was one of only eight lncRNAs identified as associated with a high-risk of gastric cancer (Zhao *et al*., 2022), but the dataset analysed in this study contained both male and female samples, meaning the prognostic value of *RP11-115H13.1* in male patients was likely underestimated. To our knowledge, there are no existing reports of the potential cellular function of *RP11-115H13.1* or *ARSFP1. TBL1Y* has been reported as involved in syndromic hearing loss (Di Stazio *et al*., 2019) and cardiac differentiation (Meyfour *et al*., 2017), but studies of its function in the stomach are lacking.

There are limitations in our study. We do not profile cell subtypes; such as those included under the umbrella term of ’gastric enteroendocrine cells’ including D-cells and G-cells, for which it was not possible to identify *Ref.T*. that fulfilled the required criteria. Our observations are consistent with these sub-cell types being typically defined by the expression of a limited number of specialised proteins (Sjölund *et al*., 1983; Engelstoft *et al*., 2013; Gribble and Reimann, 2016), rather than large distinct gene signature panels. Gene expression in stomach can be modified by genetic or environmental factors, such as the individual variation in the gastrointestinal microbiome (Nichols and Davenport, 2021). Strongly regulated genes may therefore not correlate with the more constitutively expressed *Ref.T*. selected to represent the cell type in which they are primarily expressed, as variation across samples could be independent of cell type proportions. Thus, such genes could be false negatives in our analysis. Furthermore, we have used high thresholds for the classification of genes as cell type-enriched, which could lead to incorrect exclusion. For example, tissue profiling showed that proteins encoded by *MUC4* and *MUC5B* are selectively expressed in gastric mucous cells (Uhlen *et al*., 2019), but they fall just below the threshold for classification as such in our analysis. However, the individual enrichment scores clearly indicate that they have a cell-type restricted expression; thus, our classifications should be considered only a guide, and the data should be considered on a transcript-by-transcript basis.

## ACKNOWLEDGEMENTS

Funding granted to LMB from Hjärt Lungfonden (20170759, 20170537, 20200544) and Swedish Research Council (2019-01493), and to JO from Stockholm County Council (SLL 2017-0842). The Human Protein Atlas is funded by The Knut and Alice Wallenberg Foundation. **Data usage:** We used data from Genotype-Tissue Expression (GTEx) Project (gtexportal.org) (Consortium, 2015) supported by the Office of the Director of the National Institutes of Health, and by NCI, NHGRI, NHLBI, NIDA, NIMH, and NINDS.

## AUTHOR CONTRIBUTIONS

Conceptualisation: LMB. Methodology: SÖ, ES, MNT, PD. Formal analysis: SÖ, PD, LMB. Investigation SÖ, PD, LMB, CL. Resources: MU, FP, JO, LB, CL. Writing – Original Draft: SÖ, LMB. Writing – Review & Editing: All, Visualisation: SÖ, LMB, PD, MZ, KVF. Supervision: LMB, PD. Funding Acquisition: LMB, JO.

## METHODS AND RESOURCES LEAD CONTACT

Further information and requests for resources and reagents should be directed to and will be fulfilled by the Lead Contact: Dr. Lynn Marie Butler.

## MATERIALS AVALIBILITY

This study did not generate new unique reagents.

## DATA AND CODE AVAILABILITY

- This paper analyses existing, publicly available data from the Genotype-Tissue Expression(GTEx) project with accession number phs000424.v8.p2 (Consortium, 2015) and single cell RNAseq data from Tabula Sapiens (Tabula Sapiens *et al*., 2022) retrieved on 2022/07/29.
- All original code has been deposited at GitHub and is publicly available as of the date of publication, link: https://github.com/PhilipDusart/cell-enrichment.
- Any additional information required to reanalyse the data reported in this paper is available from the lead contact upon request.

## EXPERIMENTAL MODEL AND SUBJECT DETAILS

Bulk RNAseq data analysed in this study was obtained from the Genotype-Tissue Expression (GTEx) Project (gtexportal.org) (Consortium, 2015) accessed on 2021/04/26 (dbGaP Accession phs000424.v8.p2). Transcript types were categorised according to Biotype definitions in ENSEMBL release 102 (Yates *et al*., 2020). Human tissue protein profiling was performed in house as part of the Human Protein Atlas (HPA) project (Ponten, Jirstrom and Uhlen, 2008; Uhlen *et al*., 2015, 2017) (www.proteinatlas.org). Human stomach tissue samples were obtained from the Department of Pathology, Uppsala University Hospital, Uppsala, Sweden, as part of the Uppsala Biobank. Samples were handled in accordance with Swedish laws and regulations, with approval from the Uppsala Ethical Review Board (Uhlen et al., 2015).

## METHOD DETAILS

### Tissue Profiling: Human tissue sections

Stomach tissue sections were stained, as previously described (Ponten, Jirstrom and Uhlen, 2008; Uhlen *et al*., 2015). Briefly, formalin fixed and paraffin embedded tissue samples were sectioned, de-paraffinised in xylene, hydrated in graded alcohols and blocked for endogenous peroxidase in 0.3% hydrogen peroxide diluted in 95% ethanol. For antigen retrieval, a Decloaking chamber® (Biocare Medical, CA) was used. Slides were boiled in Citrate buffer®, pH6 (Lab Vision, CA). Primary antibodies and a dextran polymer visualization system (UltraVision LP HRP polymer®, Lab Vision) were incubated for 30 min each at room temperature and slides were developed for 10 minutes using Diaminobenzidine (Lab Vision) as the chromogen. Slides were counterstained in Mayers hematoxylin (Histolab) and scanned using Scanscope XT (Aperio). Primary antibodies, source, target and identifier are as follows: Atlas Antibodies: ACSS1 (Cat#HPA043228, RRID:AB_2678372), ATP4A (Cat#HPA076684), ATP4B (Cat#HPA045400, RRID:AB_2679314), MFSD4A (Cat#055407), SH3GL2 (Cat#HPA026685, RRID:AB_1856817), SLC9A3 (Cat#HPA036493, RRID:AB_10673353), TPCN2 (Cat#HPA027080, RRID:AB_10600917), CEBPA (Cat#HPA065037, RRID:AB_2685410), LIPF (Cat#HPA045930, RRID:AB_10959518), SPTBN2 (Cat#HPA043529, RRID:AB_2678531), BHLHA15 (Cat#HPA047834, RRID:AB_2680172), KIAA1324 (Cat#HPA029869, RRID:AB_10794320), PGC (Cat#HPA031717, RRID:AB_10670130), CAMK2B (Cat#HPA053973, RRID:AB_2682328), SLC18A1 (Cat#HPA063797, RRID:AB_2685125), MS4A8 (Cat#HPA007319, RRID:AB_1854138), NKX2-2 (Cat#HPA003468, RRID:AB_1079490), TFF2 (Cat#HPA036705, RRID:AB_2675263), VILL (Cat#HPA035675, RRID:AB_10671223), CTSE (Cat#HPA012940, RRID:AB_2668773), FER1L6 (Cat#HPA054117, RRID:AB_2682387), LGALS4 (Cat#HPA031186, RRID:AB_2673778), PLAC8 (Cat#HPA040465, RRID:AB_10794875), CCNB1 (Cat#HPA061448, RRID:AB_2684522), DLGAP5 (Cat#HPA005546, RRID:AB_1078677), TPX2 (Cat#HPA005487, RRID:AB_1858223), PECAM1 (Cat#HPA004690, RRID:AB_1078462), CD93 (Cat#HPA009300, RRID:AB_1846342), MFAP2 (Cat#HPA007354, RRID:AB_1079365), MFAP4 (Cat#HPA054097, RRID:AB_2682378) EMILIN1 (Cat#HPA002822, RRID:AB_1078738), AIF1 (Cat#HPA049234, RRID:AB_2680685), ITGB2 (Cat#HPA016894, RRID:AB_1846257), CXCR2 (Cat#HPA032017, RRID:AB_2674112), PADI4 (Cat#HPA017007, RRID:AB_1854921), S100A12 (Cat#HPA002881, RRID:AB_1848175), CD2 (Cat#HPA003883, RRID:AB_1846263), CD3E (Cat#HPA043955, RRID:AB_2678747), IGHA1 (Cat#HPA001217, RRID:AB_1079120), JCHAIN (Cat#HPA044132, RRID:AB_2678826) and MZB1 (Cat#HPA043745, RRID:AB_10960359) from Santa Cruz Biotechnology: AZGP1 (Cat#sc-13585, RRID:AB_667849), PAX6 (Cat#sc-81649, RRID:AB_1127044), BIRC5 (Cat#sc-17779, RRID:AB_628302), CDC20 (Cat#sc-13162, RRID:AB_628089), S1PR1 (Cat#sc-48356, RRID:AB_2238920), FCGR3A (Cat#sc-20052, RRID:AB_626925) from Agilent: CD8A (Cat#M7103, RRID:AB_2075537) from Leica Biosystems: TOP2A (Cat#NCL-TOPOIIA, RRID:AB_564035), TFF1 (Cat#NCL-pS2, RRID:AB_563985) from Epitomics an AbCam company: CDK1 (Cat#1161-1, RRID:AB_344898) and from Roche: CHGA (Product name: 1199 021).

## QUANTIFICATION AND STATISTICAL ANALYSIS

### Reference transcript-based correlation analysis

This method was adapted and expanded from that previously developed to determine the cross-tissue pan-EC-enriched transcriptome (Butler *et al*., 2016) and human brain and adipose tissue cell-enriched genes (Dusart *et al*., 2019; Norreen-Thorsen *et al*., 2022). Pairwise Spearman correlation coefficients were calculated between reference transcripts selected as proxy markers for: parietal cells [*ATP4B, MFSD4A, ATP4A*], chief cells [*PGC, LIPF, AZGP1*], gastric enteroendocrine cells [*ST18, INSM1, ARX*], gastric mucous cells [*LGALS4, VILL, CAPN8*], mitotic cells [*NCAPG, KIFC1, NCAPH*], endothelial cells [PECAM1, CDH5, ERG], fibroblast [*PCOLCE, CLEC11A, MMP2*], macrophages [*C1QB, FCGR3A, ITGB2*], neutrophils [*CXCR2, FCGR3B, CXCR1*], T-cells [*CD3E, CD2, CD3G*] and plasma cells [*IGKC, JCHAIN, IGLC1*] and all other sequenced transcripts. Transcripts with a TPM value <0.1 in more than 50% of samples were excluded from analysis (but are still included in data tables). See results section for full criteria required for transcript classification of transcripts as cell-type enriched. Correlation coefficients were calculated in R using the *corr.test* function from the *psych* package (v 1.8.4). In addition to correlation coefficients False Discovery Rate (FDR) adjusted p-values (using Bonferroni correction) and raw p-values were calculated. FDR <0.0001 for correlation was required for inclusion as cell type enriched, but no transcripts required exclusion due to this criterion.

### Weighted correlation network (WGCNA) analysis

The R package WGCNA (Langfelder and Horvath, 2008) was used to perform co-expression network analysis for gene clustering, on log2 expression TPM values. The analysis was performed according to recommendations in the WGCNA manual. Transcripts with too many missing values were excluded using the goodSamplesGenes() function. The remaining genes were used to cluster the samples, and obvious outlier samples were excluded.

### Gene ontology and reactome analysis

The Gene Ontology Consortium (Ashburner *et al*., 2000) and PANTHER classification resource (Mi *et al*., 2013, 2016) were used to identify over represented terms (biological processes) in the panel of identified cell-type-enriched transcripts from the GO ontology (release date 2022-03-22) or reactome (release date 2021-10-01) databases. Plots of GO terms were created using REVIGO (Supek et al., 2011) where stated.

### Visualisation

Circular graphs were constructed using the R package *circlize* (Gu *et al*., 2014). Some figure sections were created with BioRender.com.

### Additional datasets and analysis

Single cell RNAseq data from Tabula Sapiens (Tabula Sapiens et al., 2022) was downloaded and UMAP plots created using the Seurat package in R (Hao et al., 2021). Tissue enriched genes were downloaded from the Human Protein Atlas (HPA) tissue atlas (Uhlen et al., 2015) or GTEx database (Consortium, 2015), as collated in the Harminozome database (Rouillard et al., 2016).

## ADDITIONAL RESOURCES

Analysed data for all protein coding genes is provided on the Human Protein Atlas website: (https://www.proteinatlas.org/humanproteome/tissue+cell+type/stomach). Data for non-coding transcripts is provided on: https://cell-enrichment.shinyapps.io/noncoding_stomach/. The published article includes all datasets generated during this study (Tables S1 and S2).

## SUPPLEMENTAL TABLE LEGENDS

***Table S1. Reference transcript selection and analysis criteria***.

(Tab **1)**: Correlation coefficient values were calculated between selected *Ref.T*. to represent constituent stomach cell types. (Tab **2)**: Correlation coefficient values were calculated between selected *Ref.T*. and all other sequenced transcripts in GTEx stomach mRNAseq data (Table A) and the mean differential vs. all *Ref.T*. panels (Table B). Transcripts classified as enriched in: (Tab **3**) parietal cells, (Tab **4**) chief cells, (Tab **5**) gastric enteroendocrine cells, (Tab **6**) gastric mucous cells, (Tab **7**) mitotic cells, (Tab **8**) endothelial cells, (Tab **9**) fibroblasts, (Tab **10**) macrophages, (Tab **11**) neutrophils, (Tab **12**) T-cells and (Tab **13**) plasma cells were analysed to identify over-represented terms in the (Table **A**) gene ontology or (Table **B**). *Related to all Figures*.

**Table S2. Sex stratified subset analysis of cell-enriched transcripts in human stomach**. (Tab **1)**: Correlation coefficient values were calculated between selected *Ref.T*. to represent constituent stomach cell types in females (Table **A**) or males (Table **B**). (Tab **2**) Correlation coefficient values were calculated between selected *Ref.T*. and all other sequenced transcripts in stomach mRNAseq data (GTEx), subdivided into (Table **A**) female or (Table **B**) male only sample sets. See key for column details. *Related to Figure 7 and S3*.

---

**Figure S1.**
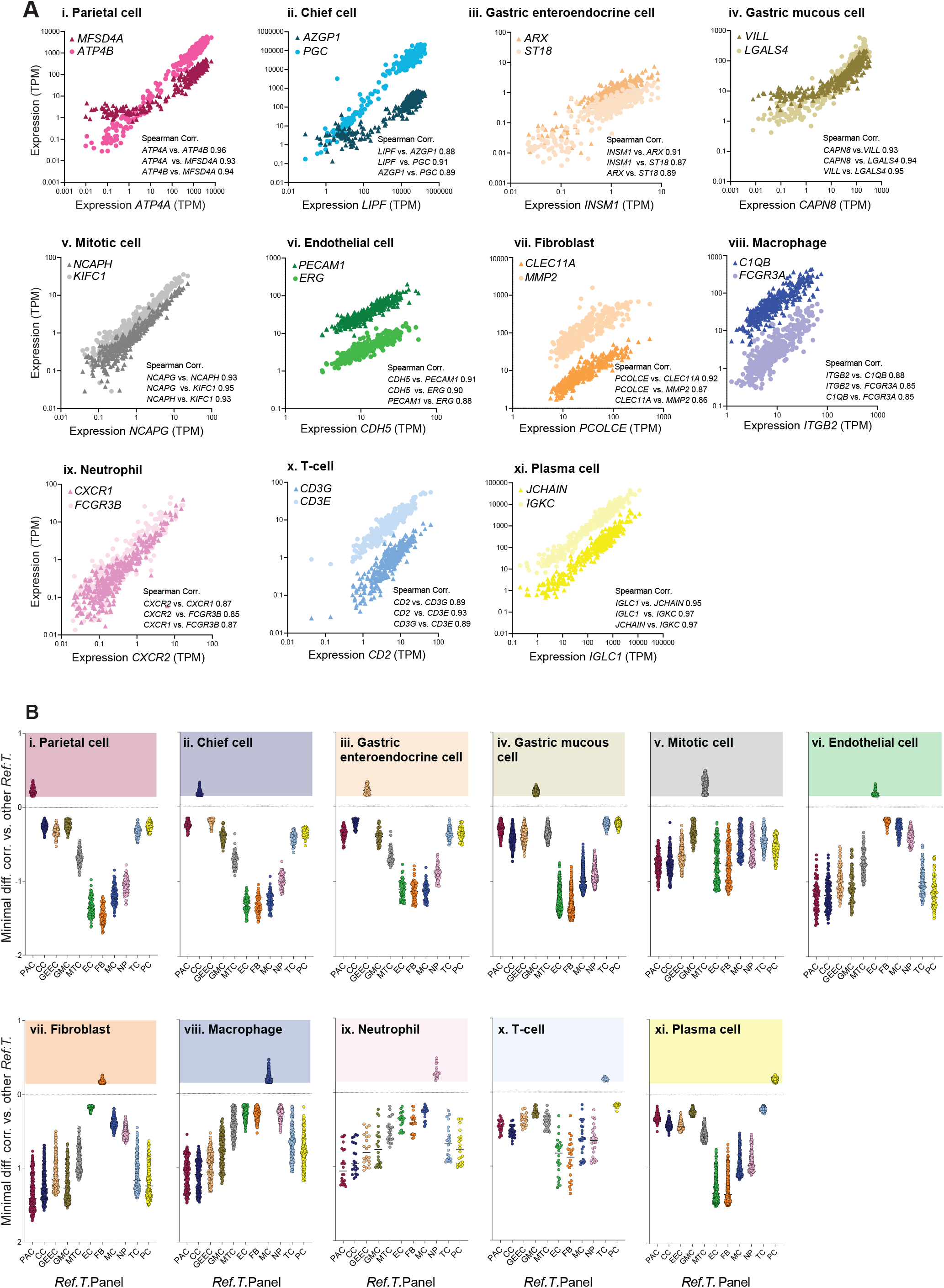
Expression distribution and correlations between human stomach cell type reference transcripts; Related to Figure 1 and Table S1, Tab 1. (**A**) Expression of *Ref.T* selected to represent: (i) parietal cells, (ii) chief cells, (iii) gastric enteroendocrine cells, (iv) gastric mucous cells, (v) mitotic cells, (vi) endothelial cells, (vii) fibroblasts, (viii) macrophages, (ix) neutrophils, (x) T-cells and (xi) plasma cells. (**B**) Minimal differential correlations between mean correlation coefficients with corresponding *Ref.T*. panel for genes above designated thresholds for classification as cell type enriched in: (i) parietal cells, (ii) chief cells, (iii) gastric enteroendocrine cells, (iv) gastric mucous cells, (v) mitotic cells, (vi) endothelial cells, (vii) fibroblasts, (viii) macrophages, (ix) neutrophils, (x) T-cells and (xi) plasma cells, and the mean correlation coefficients with all other *Ref.T*. panels.

**Figure S2.**
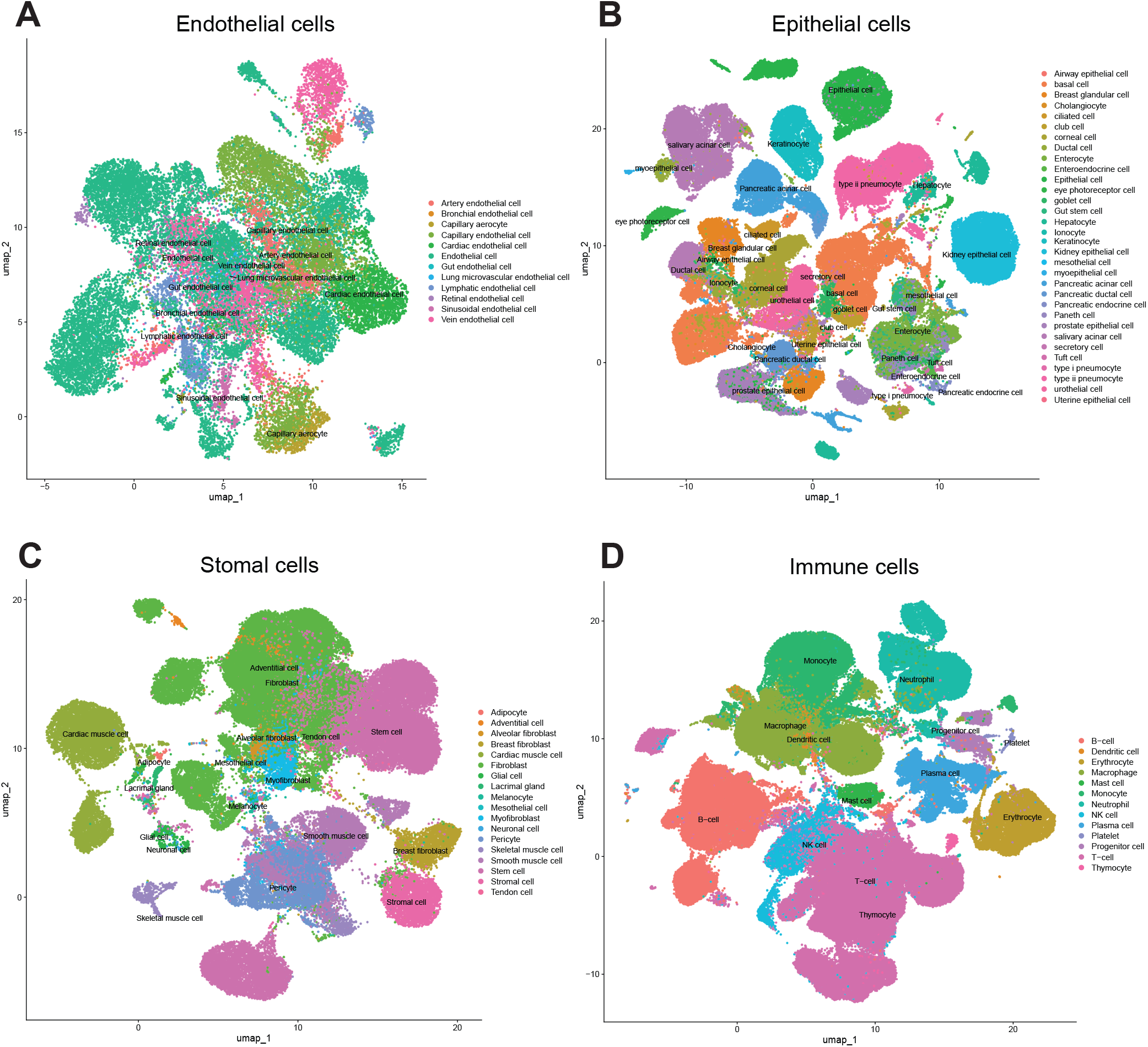
Single cell RNAseq (scRNAseq) annotations; Related to Figure 5, 6 and 7. scRNAseq data was sourced from Tabula Sapiens (Tabula Sapiens et al., 2022). UMAP plots showing original annotations of cell clusters designated as: (**A**) endothelial, (**B**) epithelial, (**C**) stromal or (**D**) immune.

**Figure S3.**
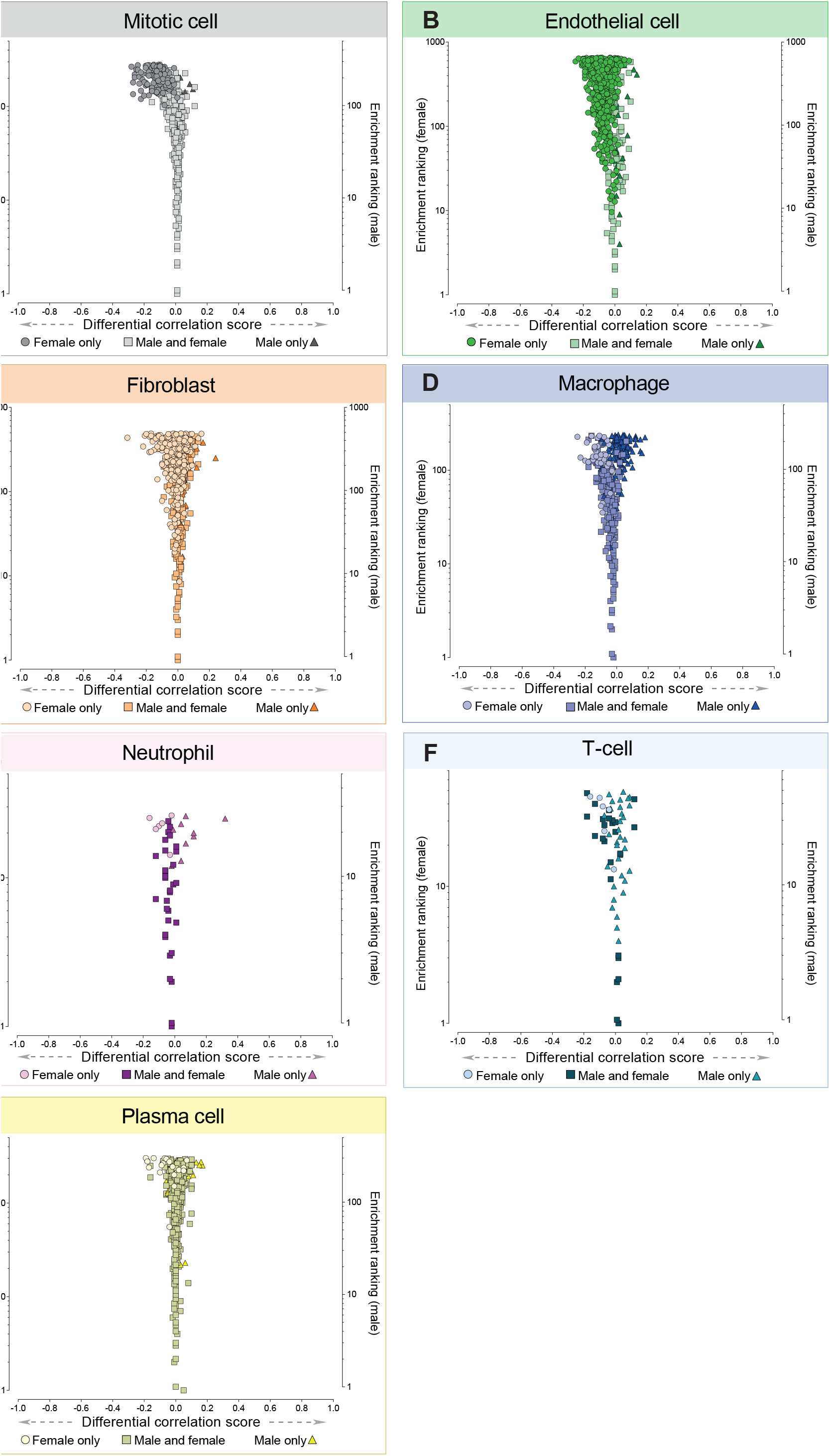
Identification of sex-specific cell type enriched genes in human stomach; Related to Figure 7. Human stomach RNAseq data (n=359 individuals) was retrieved from GTEx V8 and divided into female (n=132) and male (n=227) subgroups before classification of cell type-enriched transcripts. For genes classified as: (**A**) mitotic cell, (**B**) endothelial cell, (**C**) fibroblast, (**D**) macrophage, (**E**) neutrophil, (**F**) T-cell and (**G**) plasma cell enriched, in either female or male subsets, the ’*sex differential correlation score*’ (difference between mean correlation with the *Ref.T* panel in females vs. males) was plotted vs. ‘enrichment ranking’ (position in each respective enriched list, highest correlation = rank 1). See also Table S2, Tab 1. On each plot, transcripts enriched in *both* females and males are represented by common-coloured square symbols, and transcripts classified as enriched *only* in females or males are represented by differently coloured circle and triangle symbols, respectively. See also Table S2, Tab 1.

## REFERENCES

Aihara, E., Engevik, K.A. and Montrose, M.H. (2017) ‘Trefoil Factor Peptides and Gastrointestinal Function’, Annual Review of Physiology, 79, pp. 357–380. Available at: https://doi.org/10.1146/annurev-physiol-021115-105447.

Alpers, D.H. and Russell-Jones, G. (2013) ‘Gastric intrinsic factor: The gastric and small intestinal stages of cobalamin absorption. A personal journey’, Biochimie, 95(5), pp. 989–994. Available at: https://doi.org/10.1016/j.biochi.2012.12.006.

Al-Shboul, O. (2016) ‘The role of the RhoA/ROCK pathway in gender-dependent differences in gastric smooth muscle contraction’, The Journal of Physiological Sciences, 66(1), pp. 85–92. Available at: https://doi.org/10.1007/s12576-015-0400-9.

Ashburner, M. et al. (2000) ‘Gene Ontology: tool for the unification of biology’, Nature Genetics, 25(1), pp. 25–29. Available at: https://doi.org/10.1038/75556.

Beucher, A. et al. (2012) ‘The Homeodomain-Containing Transcription Factors Arx and Pax4 Control Enteroendocrine Subtype Specification in Mice’, PLOS ONE, 7(5), p. e36449. Available at: https://doi.org/10.1371/journal.pone.0036449.

Busslinger, G.A. et al. (2021) ‘Human gastrointestinal epithelia of the esophagus, stomach, and duodenum resolved at single-cell resolution’, Cell Reports, 34(10), p. 108819. Available at: https://doi.org/10.1016/j.celrep.2021.108819.

Butler, L.M. et al. (2016) ‘Analysis of Body-wide Unfractionated Tissue Data to Identify a Core Human Endothelial Transcriptome’, Cell Systems, 3(3), pp. 287-301.e3. Available at: https://doi.org/10.1016/j.cels.2016.08.001.

Cai, L. et al. (2007) ‘Identification of PRTFDC1 silencing and aberrant promoter methylation of GPR150, ITGA8 and HOXD11 in ovarian cancers’, Life Sciences, 80(16), pp. 1458–1465. Available at: https://doi.org/10.1016/j.lfs.2007.01.015.

Capizzi, M. et al. (2017) ‘MIR7-3HG, a MYC-dependent modulator of cell proliferation, inhibits autophagy by a regulatory loop involving AMBRA1’, Autophagy, 13(3), pp. 554–566. Available at: https://doi.org/10.1080/15548627.2016.1269989.

Cheetham, S.W., Faulkner, G.J. and Dinger, M.E. (2020) ‘Overcoming challenges and dogmas to understand the functions of pseudogenes’, Nature Reviews Genetics, 21(3), pp. 191–201. Available at: https://doi.org/10.1038/s41576-019-0196-1.

Cho, C.J., Park, D. and Mills, J.C. (2022) ‘ELAPOR1 is a secretory granule maturation-promoting factor that is lost during paligenosis’, American Journal of Physiology-Gastrointestinal and Liver Physiology, 322(1), pp. G49–G65. Available at: https://doi.org/10.1152/ajpgi.00246.2021.

Choi, E. et al. (2014) ‘Cell lineage distribution atlas of the human stomach reveals heterogeneous gland populations in the gastric antrum’, Gut, 63(11), pp. 1711–1720. Available at: https://doi.org/10.1136/gutjnl-2013-305964.

Choi, W.S. et al. (2013) ‘Gastrokine 1 expression in the human gastric mucosa is closely associated with the degree of gastritis and DNA methylation’, Journal of Gastric Cancer, 13(4), pp. 232–241. Available at: https://doi.org/10.5230/jgc.2013.13.4.232.

Consortium, G.Te. (2015) ‘Human genomics. The Genotype-Tissue Expression (GTEx) pilot analysis: multitissue gene regulation in humans’, Science, 348(6235), pp. 648–60. Available at: https://doi.org/10.1126/science.1262110.

Datz, F.L., Christian, P.E. and Moore, J. (1987) ‘Gender-related differences in gastric emptying’, Journal of Nuclear Medicine: Official Publication, Society of Nuclear Medicine, 28(7), pp. 1204–1207.

Denisenko, E. et al. (2020) ‘Systematic assessment of tissue dissociation and storage biases in single-cell and single-nucleus RNA-seq workflows’, Genome Biol. 2020/06/04 edn, 21(1), p. 130. Available at: https://doi.org/10.1186/s13059-020-02048-6.

Denninger, J.K. et al. (2022) ‘Robust Transcriptional Profiling and Identification of Differentially Expressed Genes With Low Input RNA Sequencing of Adult Hippocampal Neural Stem and Progenitor Populations’, Frontiers in Molecular Neuroscience, 15, p. 810722. Available at: https://doi.org/10.3389/fnmol.2022.810722.

Di Stazio, M. et al. (2019) ‘TBL1Y: a new gene involved in syndromic hearing loss’, European Journal of Human Genetics, 27(3), pp. 466–474. Available at: https://doi.org/10.1038/s41431-018-0282-4.

Dusart, P. et al. (2019) ‘A Systems-Based Map of Human Brain Cell-Type Enriched Genes and Malignancy-Associated Endothelial Changes’, Cell Reports, 29(6), pp. 1690-1706.e4. Available at: https://doi.org/10.1016/j.celrep.2019.09.088.

Engelstoft, M.S. et al. (2013) ‘Enteroendocrine cell types revisited’, Current Opinion in Pharmacology, 13(6), pp. 912–921. Available at: https://doi.org/10.1016/j.coph.2013.09.018.

Franzen, O., Gan, L.M. and Bjorkegren, J.L.M. (2019) ‘PanglaoDB: a web server for exploration of mouse and human single-cell RNA sequencing data’, Database (Oxford). 2019/04/06 edn, 2019. Available at: https://doi.org/10.1093/database/baz046.

Fujita, Y. et al. (2008) ‘Pax6 and Pdx1 are required for production of glucose-dependent insulinotropic polypeptide in proglucagon-expressing L cells’, American Journal of Physiology-Endocrinology and Metabolism, 295(3), pp. E648–E657. Available at: https://doi.org/10.1152/ajpendo.90440.2008.

Gao, Y. et al. (2020) ‘Long noncoding RNAs in gastric cancer: From molecular dissection to clinical application’, World J Gastroenterol. 2020/07/14 edn, 26(24), pp. 3401–3412. Available at: https://doi.org/10.3748/wjg.v26.i24.3401.

Gawad, C., Koh, W. and Quake, S.R. (2016) ‘Single-cell genome sequencing: current state of the science’, Nature Reviews Genetics, 17(3), pp. 175–188. Available at: https://doi.org/10.1038/nrg.2015.16.

Gene Ontology, C. (2021) ‘The Gene Ontology resource: enriching a GOld mine’, Nucleic Acids Res. 2020/12/09 edn, 49(D1), pp. D325–D334. Available at: https://doi.org/10.1093/nar/gkaa1113.

Ghafouri-Fard, S. and Taheri, M. (2020) ‘Long non-coding RNA signature in gastric cancer’, Experimental and Molecular Pathology, 113, p. 104365. Available at: https://doi.org/10.1016/j.yexmp.2019.104365.

Goldspink, D.A., Reimann, F. and Gribble, F.M. (2018) ‘Models and Tools for Studying Enteroendocrine Cells’, Endocrinology, 159(12), pp. 3874–3884. Available at: https://doi.org/10.1210/en.2018-00672.

Gremel, G. et al. (2015) ‘The human gastrointestinal tract-specific transcriptome and proteome as defined by RNA sequencing and antibody-based profiling’, Journal of Gastroenterology, 50(1), pp. 46–57. Available at: https://doi.org/10.1007/s00535-014-0958-7.

Gribble, F.M. and Reimann, F. (2016) ‘Enteroendocrine Cells: Chemosensors in the Intestinal Epithelium’, Annual Review of Physiology, 78(1), pp. 277–299. Available at: https://doi.org/10.1146/annurev-physiol-021115-105439.

Grün, D. and van Oudenaarden, A. (2015) ‘Design and Analysis of Single-Cell Sequencing Experiments’, Cell, 163(4), pp. 799–810. Available at: https://doi.org/10.1016/j.cell.2015.10.039.

Gu, Z. et al. (2014) ‘circlize Implements and enhances circular visualization in R’, Bioinformatics. 2014/06/16 edn, 30(19), pp. 2811–2. Available at: https://doi.org/10.1093/bioinformatics/btu393.

Hanasaki, K. et al. (2002) ‘Potent Modification of Low Density Lipoprotein by Group X Secretory Phospholipase A2 Is Linked to Macrophage Foam Cell Formation *’, Journal of Biological Chemistry, 277(32), pp. 29116–29124. Available at: https://doi.org/10.1074/jbc.M202867200.

Hassan, M.I., Toor, A. and Ahmad, F. (2010) ‘Progastriscin: structure, function, and its role in tumor progression’, J Mol Cell Biol. 2010/03/17 edn, 2(3), pp. 118–27. Available at: https://doi.org/10.1093/jmcb/mjq001.

Hata, S. et al. (2010) ‘Calpain 8/nCL-2 and calpain 9/nCL-4 constitute an active protease complex, G-calpain, involved in gastric mucosal defense’, PLoS genetics, 6(7), p. e1001040. Available at: https://doi.org/10.1371/journal.pgen.1001040.

Hill, M.E., Asa, S.L. and Drucker, D.J. (1999) ‘Essential Requirement for Pax6 in Control of Enteroendocrine Proglucagon Gene Transcription’, Molecular Endocrinology, 13(9), pp. 1474–1486. Available at: https://doi.org/10.1210/mend.13.9.0340.

Hooks, S.B., Ragan, S.P. and Lynch, K.R. (1998) ‘Identification of a novel human phosphatidic acid phosphatase type 2 isoform’, FEBS Letters, 427(2), pp. 188–192. Available at: https://doi.org/10.1016/S0014-5793(98)00421-9.

Ja, G. et al. (2020) ‘Mucins in Intestinal Mucosal Defense and Inflammation: Learning From Clinical and Experimental Studies’, Frontiers in immunology, 11. Available at: https://doi.org/10.3389/fimmu.2020.02054.

Jiang, R. et al. (2022) ‘Statistics or biology: the zero-inflation controversy about scRNA-seq data’, Genome Biol. 2022/01/23 edn, 23(1), p. 31. Available at: https://doi.org/10.1186/s13059-022-02601-5.

Jiang, S., Tan, B. and Zhang, X. (2019) ‘Identification of key lncRNAs in the carcinogenesis and progression of colon adenocarcinoma by co-expression network analysis’, Journal of Cellular Biochemistry, 120(4), pp. 6490–6501. Available at: https://doi.org/10.1002/jcb.27940.

Kang, Y. et al. (2015) ‘PPARG Modulated Lipid Accumulation in Dairy GMEC via Regulation of ADRP Gene’, Journal of Cellular Biochemistry, 116(1), pp. 192–201. Available at: https://doi.org/10.1002/jcb.24958.

Karlsson, M. et al. (2021) ‘A single-cell type transcriptomics map of human tissues’, Sci Adv. 2021/07/30 edn, 7(31). Available at: https://doi.org/10.1126/sciadv.abh2169.

Kim, J. et al. (2022) ‘Single-cell analysis of gastric pre-cancerous and cancer lesions reveals cell lineage diversity and intratumoral heterogeneity’, NPJ Precis Oncol. 2022/01/29 edn, 6(1), p. 9. Available at: https://doi.org/10.1038/s41698-022-00251-1.

Kim, T.H. and Shivdasani, R.A. (2016) ‘Stomach development, stem cells and disease’, Development. 2016/02/18 edn, 143(4), pp. 554–65. Available at: https://doi.org/10.1242/dev.124891.

Kirsch, S. et al. (2004) ‘Molecular and evolutionary analysis of the growth-controlling region on the human Y chromosome’, Human Genetics, 114(2), pp. 173–181. Available at: https://doi.org/10.1007/s00439-003-1028-z.

Kovalenko, T.F. and Patrushev, L.I. (2018) ‘Pseudogenes as Functionally Significant Elements of the Genome’, Biochemistry (Moscow), 83(11), pp. 1332–1349. Available at: https://doi.org/10.1134/S0006297918110044.

Langfelder, P. and Horvath, S. (2008) ‘WGCNA: an R package for weighted correlation network analysis’, BMC Bioinformatics, 9(1), p. 559. Available at: https://doi.org/10.1186/1471-2105-9-559.

Leja, J. et al. (2009) ‘Novel markers for enterochromaffin cells and gastrointestinal neuroendocrine carcinomas’, Modern Pathology: An Official Journal of the United States and Canadian Academy of Pathology, Inc, 22(2), pp. 261–272. Available at: https://doi.org/10.1038/modpathol.2008.174.

Lennerz, J.K.M. et al. (2010) ‘The Transcription Factor MIST1 Is a Novel Human Gastric Chief Cell Marker Whose Expression Is Lost in Metaplasia, Dysplasia, and Carcinoma’, The American Journal of Pathology, 177(3), pp. 1514–1533. Available at: https://doi.org/10.2353/ajpath.2010.100328.

Li, H. et al. (2020) ‘Gender Differences in Gastric Cancer Survival: 99,922 Cases Based on the SEER Database’, Journal of Gastrointestinal Surgery, 24(8), pp. 1747–1757. Available at: https://doi.org/10.1007/s11605-019-04304-y.

Li, P.-F. et al. (2014) ‘Non-coding RNAs and gastric cancer’, World Journal of Gastroenterology□: WJG, 20(18), pp. 5411–5419. Available at: https://doi.org/10.3748/wjg.v20.i18.5411.

Lou, L. et al. (2020) ‘Sex difference in incidence of gastric cancer: an international comparative study based on the Global Burden of Disease Study 2017’, BMJ open, 10(1), p. e033323. Available at: https://doi.org/10.1136/bmjopen-2019-033323.

Massoni-Badosa, R. et al. (2020) ‘Sampling time-dependent artifacts in single-cell genomics studies’, Genome Biol. 2020/05/13 edn, 21(1), p. 112. Available at: https://doi.org/10.1186/s13059-020-02032-0.

Meyfour, A. et al. (2017) ‘Y Chromosome Missing Protein, TBL1Y, May Play an Important Role in Cardiac Differentiation’, Journal of Proteome Research, 16(12), pp. 4391–4402. Available at: https://doi.org/10.1021/acs.jproteome.7b00391.

Mi, H. et al. (2013) ‘Large-scale gene function analysis with the PANTHER classification system’, Nat Protoc, 8(8), pp. 1551–66. Available at: https://doi.org/10.1038/nprot.2013.092.

Mi, H. et al. (2016) ‘PANTHER version 10: expanded protein families and functions, and analysis tools’, Nucleic Acids Res, 44(D1), pp. D336–42. Available at: https://doi.org/10.1093/nar/gkv1194.

Nichols, R.G. and Davenport, E.R. (2021) ‘The relationship between the gut microbiome and host gene expression: a review’, Human Genetics, 140(5), pp. 747–760. Available at: https://doi.org/10.1007/s00439-020-02237-0.

Norreen-Thorsen, M. et al. (2022) ‘A human adipose tissue cell-type transcriptome atlas’, Cell Reports, 40(2), p. 111046. Available at: https://doi.org/10.1016/j.celrep.2022.111046.

O’Flanagan, C.H. et al. (2019) ‘Dissociation of solid tumor tissues with cold active protease for single-cell RNA-seq minimizes conserved collagenase-associated stress responses’, Genome Biol. 2019/10/19 edn, 20(1), p. 210. Available at: https://doi.org/10.1186/s13059-019-1830-0.

Petrovic, S. et al. (2003) ‘Identification of a basolateral Cl−/HCO 3 − exchanger specific to gastric parietal cells’, American Journal of Physiology-Gastrointestinal and Liver Physiology, 284(6), pp. G1093–G1103. Available at: https://doi.org/10.1152/ajpgi.00454.2002.

Pillai, A. et al. (2007) ‘Lhx1 and Lhx5 maintain the inhibitory-neurotransmitter status of interneurons in the dorsal spinal cord’, Development, 134(2), pp. 357–366. Available at: https://doi.org/10.1242/dev.02717.

Pink, R.C. et al. (2011) ‘Pseudogenes: Pseudo-functional or key regulators in health and disease?’, RNA, 17(5), pp. 792–798. Available at: https://doi.org/10.1261/rna.2658311.

Ponten, F., Jirstrom, K. and Uhlen, M. (2008) ‘The Human Protein Atlas - a tool for pathology’, Journal of Pathology, 216(4), pp. 387–393. Available at: https://doi.org/10.1002/path.2440.

Razavi, H. and Katanforosh, A. (2022) ‘Identification of novel key regulatory lncRNAs in gastric adenocarcinoma’, BMC Genomics. 2022/05/08 edn, 23(1), p. 352. Available at: https://doi.org/10.1186/s12864-022-08578-6.

Regev, A. et al./person-group>. (2017) ‘The Human Cell Atlas’, eLife. edited by T.R. Gingeras, 6, p. e27041. Available at: https://doi.org/10.7554/eLife.27041.

Rouillard, A.D. et al. (2016) ‘The harmonizome: a collection of processed datasets gathered to serve and mine knowledge about genes and proteins’, Database (Oxford). 2016/07/05 edn, 2016. Available at: https://doi.org/10.1093/database/baw100.

de Santa Barbara, P., van den Brink, G.R. and Roberts, D.J. (2003) ‘Development and differentiation of the intestinal epithelium’, Cellular and Molecular Life Sciences (CMLS), 60(7), pp. 1322–1332. Available at: https://doi.org/10.1007/s00018-003-2289-3.

Sathe, A. et al. (2020) ‘Single-Cell Genomic Characterization Reveals the Cellular Reprogramming of the Gastric Tumor Microenvironment’, Clinical Cancer Research, 26(11), pp. 2640–2653. Available at: https://doi.org/10.1158/1078-0432.CCR-19-3231.

Schwer, B. et al. (2006) ‘Reversible lysine acetylation controls the activity of the mitochondrial enzyme acetyl-CoA synthetase 2’, Proceedings of the National Academy of Sciences, 103(27), pp. 10224–10229. Available at: https://doi.org/10.1073/pnas.0603968103.

Shapiro, E., Biezuner, T. and Linnarsson, S. (2013) ‘Single-cell sequencing-based technologies will revolutionize whole-organism science’, Nature Reviews Genetics, 14(9), pp. 618–630. Available at: https://doi.org/10.1038/nrg3542.

Shimizu, D., Kanda, M. and Kodera, Y. (2018) ‘Emerging evidence of the molecular landscape specific for hematogenous metastasis from gastric cancer’, World Journal of Gastrointestinal Oncology, 10(6), pp. 124–136. Available at: https://doi.org/10.4251/wjgo.v10.i6.124.

Sjölund, K. et al. (1983) ‘Endocrine Cells in Human Intestine: An Immunocytochemical Study’, Gastroenterology, 85(5), pp. 1120–1130. Available at: https://doi.org/10.1016/S0016-5085(83)80080-8.

Squair, J.W. et al. (2021) ‘Confronting false discoveries in single-cell differential expression’, Nature Communications, 12(1), p. 5692. Available at: https://doi.org/10.1038/s41467-021-25960-2.

Tabula Sapiens, C. et al. (2022) ‘The Tabula Sapiens: A multiple-organ, single-cell transcriptomic atlas of humans’, Science. 2022/05/14 edn, 376(6594), p. eabl4896. Available at: https://doi.org/10.1126/science.abl4896.

Thompson, C.A., DeLaForest, A. and Battle, M.A. (2018) ‘Patterning the gastrointestinal epithelium to confer regional-specific functions’, Developmental Biology, 435(2), pp. 97–108. Available at: https://doi.org/10.1016/j.ydbio.2018.01.006.

Toulza, E. et al. (2007) ‘Large-scale identification of human genes implicated in epidermal barrier function’, Genome Biology, 8(6), p. R107. Available at: https://doi.org/10.1186/gb-2007-8-6-r107.

Tsakmaki, A. et al. (2020) ‘ISX-9 manipulates endocrine progenitor fate revealing conserved intestinal lineages in mouse and human organoids’, Molecular Metabolism, 34, pp. 157–173. Available at: https://doi.org/10.1016/j.molmet.2020.01.012.

Tsubosaka, A. et al. (2022) ‘Single-Cell Transcriptome Analyses Reveal the Cell Diversity and Developmental Features of Human Gastric and Metaplastic Mucosa’. bioRxiv, p. 2022.05.22.493006. Available at: https://doi.org/10.1101/2022.05.22.493006.

Uhlen, M. et al. (2015) ‘Proteomics. Tissue-based map of the human proteome’, Science. 2015/01/24 edn, 347(6220), p. 1260419. Available at: https://doi.org/10.1126/science.1260419.

Uhlen, M. et al. (2017) ‘A pathology atlas of the human cancer transcriptome’, Science, 357(6352). Available at: https://doi.org/10.1126/science.aan2507.

Uhlen, M. et al. (2019) ‘A genome-wide transcriptomic analysis of protein-coding genes in human blood cells’, Science, 366(6472), p. eaax9198. Available at: https://doi.org/10.1126/science.aax9198.

Wang, P. et al. (2018) ‘A Novel LncRNA-miRNA-mRNA Triple Network Identifies LncRNA RP11-363E7.4 as An Important Regulator of miRNA and Gene Expression in Gastric Cancer’, Cellular Physiology and Biochemistry, 47(3), pp. 1025–1041. Available at: https://doi.org/10.1159/000490168.

Wang, R. et al. (2021) ‘Single-cell dissection of intratumoral heterogeneity and lineage diversity in metastatic gastric adenocarcinoma’, Nature Medicine, 27(1), pp. 141–151. Available at: https://doi.org/10.1038/s41591-020-1125-8.

Wei, L. et al. (2020) ‘Noncoding RNAs in gastric cancer: implications for drug resistance’, Molecular Cancer, 19(1), p. 62. Available at: https://doi.org/10.1186/s12943-020-01185-7.

Xia, T. et al. (2015) ‘Long noncoding RNA FER1L4 suppresses cancer cell growth by acting as a competing endogenous RNA and regulating PTEN expression’, Scientific Reports, 5(1), p. 13445. Available at: https://doi.org/10.1038/srep13445.

Yan, H.-T. et al. (2005) ‘Molecular analysis of TBL1Y, a Y-linked homologue of TBL1X related with X-linked late-onset sensorineural deafness’, Journal of Human Genetics, 50(4), pp. 175–181. Available at: https://doi.org/10.1007/s10038-005-0237-9.

Yang, J.-K. et al. (2018) ‘From Hyper-to Hypoinsulinemia and Diabetes: Effect of KCNH6 on Insulin Secretion’, Cell Reports, 25(13), pp. 3800-3810.e6. Available at: https://doi.org/10.1016/j.celrep.2018.12.005.

Yang, X.-Z. et al. (2018) ‘LINC01133 as ceRNA inhibits gastric cancer progression by sponging miR-106a-3p to regulate APC expression and the Wnt/β-catenin pathway’, Molecular Cancer, 17(1), p. 126. Available at: https://doi.org/10.1186/s12943-018-0874-1.

Yates, A.D. et al. (2020) ‘Ensembl 2020’, Nucleic Acids Research, 48(D1), pp. D682–D688. Available at: https://doi.org/10.1093/nar/gkz966.

Yen, C.-L.E. et al. (2002) ‘Identification of a gene encoding MGAT1, a monoacylglycerol acyltransferase’, Proceedings of the National Academy of Sciences of the United States of America, 99(13), pp. 8512–8517. Available at: https://doi.org/10.1073/pnas.132274899.

Zhang, P. et al. (2019) ‘Dissecting the Single-Cell Transcriptome Network Underlying Gastric Premalignant Lesions and Early Gastric Cancer’, Cell Reports, 27(6), pp. 1934-1947.e5. Available at: https://doi.org/10.1016/j.celrep.2019.04.052.

Zhang, X. et al. (2019) ‘CellMarker: a manually curated resource of cell markers in human and mouse’, Nucleic Acids Res. 2018/10/06 edn, 47(D1), pp. D721–D728. Available at: https://doi.org/10.1093/nar/gky900.

Zhao, X. et al. (2022) ‘An Immunity-Associated lncRNA Signature for Predicting Prognosis in Gastric Adenocarcinoma’, Journal of Healthcare Engineering, 2022, p. e3035073. Available at: https://doi.org/10.1155/2022/3035073.

Zhao, Y. et al. (1999) ‘Control of Hippocampal Morphogenesis and Neuronal Differentiation by the LIM Homeobox Gene Lhx5’, Science, 284(5417), pp. 1155–1158. Available at: https://doi.org/10.1126/science.284.5417.1155.

Zheng, L. et al. (2021) ‘Long noncoding RNA LINC00982 upregulates CTSF expression to inhibit gastric cancer progression via the transcription factor HEY1’, American Journal of Physiology-Gastrointestinal and Liver Physiology, 320(5), pp. G816–G828. Available at: https://doi.org/10.1152/ajpgi.00209.2020.

